# VitTCR: A deep learning method for peptide recognition prediction

**DOI:** 10.1101/2023.06.02.543411

**Authors:** Mengnan Jiang, Zilan Yu, Xun Lan

**Affiliations:** School of Medicine, Tsinghua University, 100084, Beijing, China; Centre for Life Sciences, Tsinghua University, 100084, Beijing, China; Tsinghua-Peking Center for Life Sciences, MOE Key Laboratory of Tsinghua University, Beijing, China; MOE Key Laboratory of Bioinformatics, Tsinghua University, 100084, Beijing, China

## Abstract

The identification of the interaction between T-cell receptors (TCRs) and immunogenic peptides is important for the development of novel cancer immunotherapies and vaccines. However, experimentally determining whether a TCR recognizes a peptide is still time– and labour-consuming. In this study, we introduced VitTCR, a predictive model based on the architecture of the vision transformer (ViT), designed to forecast TCR-peptide interactions. Prior to prediction, VitTCR converts the TCR-peptide interactions into a numerical tensor named AtchleyMaps using Atchley factors. Subsequently, VitTCR takes AtchleyMaps as inputs and predicts whether an interaction between a TCR and a peptide exists. Through comprehensive evaluations, we demonstrate that VitTCR surpasses other published methods in classifying TCR-peptide pairs, exhibiting superior performance in terms of the area under the receiver operating characteristic (AUROC) and the area under the precision-recall curve (AUPR). To determine the focal contact point between TCRs and peptides, we obtained a positional bias weight matrix (PBWM) from the empirical amino acid (AA) contact probabilities derived from 83 structurally resolved pMHC-TCR complexes. The comparison of VitTCR with and without the integration of the PBWM revealed significant enhancements in the performance of the model. Moreover, the predicted probabilities generated by VitTCR exhibit significant correlations with immunological factors such as the clonal expansion and activation percentages of T cells. This further supports the efficacy of VitTCR in capturing biologically meaningful TCR-peptide interactions. In conclusion, VitTCR provides a useful computational tool for the prediction of TCR-peptide interactions, thereby contributing to our understanding in this field.

## Introduction

T-cell receptors (TCRs) are heterodimers immobilized on the surface of T cells that recognize antigenic peptides presented by the major histocompatibility complex (MHC). Ninety-five percent of T cells are composed of highly variable α and β subunits linked by disulphide bonds, which are named αβ T cells. The remaining T cells are called γδ T cells and consist of γ and δ subunits^1^. The diversity of TCRs is mainly derived from the V(D)J recombination of immunoglobulin genes, with α and γ subunits arising from VJ recombination and β and δ subunits forming from VDJ recombination. Due to somatic recombination and random insertion of nucleotides during T-cell development, humans have a highly diverse TCR repertoire, containing 1015^2^ to 1061^3^ possible receptors. The variable (V) region of the TCR subunits is responsible for the recognition of peptide-MHCs (pMHCs). The V region contains three highly variable complementarity-determining regions (CDRs): CDR1, CDR2, and CDR3. CDR3 is responsible for direct contact with antigenic peptides, which play an essential role in the recognition process. CD4 and CD8 are proteins expressed on the membrane surfaces of helper T cells and cytotoxic T cells, respectively. They can enhance the sensitivity and responses of T cells to pMHC^4,5^. This study focused on the CDR3 region of the TCRs of cytotoxic CD8+ T cells.

Previous studies have mainly focused on the binding between MHCs and antigenic peptides, with methods such as ACME^6^, NetMHCpan-4.0^7^, DeepLigand^8^, MHCflurry^9^, DeepSeqPan^10^, MHCSeqNet^11^, and DCNN^12^, all of which are trained on the affinity data between MHCs and antigenic peptides. However, effective learning of TCR-pMHC recognition remains challenging due to the lack of a training dataset and the high diversity of TCR sequences. Fortunately, with the development of sequencing technology, many more experimental datasets have been generated, including VDJdb^13^, IEDB^14^, and McPAS-TCR^15^. Several studies have shown that TCRs with similar CDR3 sequences are more likely to recognize the same peptide. While TCR classification methods, such as TCRdist^16^, GLIPH^17^, and TCRGP^18^, mainly focus on TCR sequences, interaction prediction methods, such as NetTCR2.0^19^, TITAN^20^, ImRex^21^, ERGO^22^, and pMTnet^23^, take peptide sequences into account. These models encode the AA sequences of TCRs and antigenic peptides separately and then concatenate them for feature extraction to predict whether an interaction occurs in a TCR-peptide pair.

Building a precise and robust model to predict TCR-peptide interactions remains challenging due to the high diversity of the TCR repertoire and various technological limitations. For instance, an antigenic epitope can be recognized by multiple T-cell clonotypes^16,17,24^, while a T-cell clonotype can exhibit cross-reactivity to multiple antigenic epitopes^25^. In addition, both α and β chains are considered to contribute to the binding specificity of TCR peptides^18,19^. However, despite the advent of single-cell TCR paired-strand sequencing, currently available TCR epitope binding data still mainly consist of single-strand (β-strand) information.

In this study, we leveraged the architecture of ViT^26^, initially designed for image recognition, to develop a novel model named VitTCR. VitTCR encodes the CDR3β-peptide interactions into AtchleyMaps and then takes the AtchleyMaps as inputs and outputs the predicted binding probabilities. To comprehensively assess the performance of VitTCR, we conducted comprehensive evaluations using independent testing datasets. The results demonstrated that VitTCR could be used to accurately predict whether binding occurs between MHC-I-presented epitopes and CDR3βs. Notably, comparative analyses illustrated the superior performance of VitTCR compared to other existing methods, highlighting its efficacy and potential. Moreover, the prediction of VitTCR matches the results from T-cell clonal expansions and in vitro T-cell activation experiments.

## Results

### The architecture of VitTCR

We have developed VitTCR, a deep learning-based method specifically designed to predict the interactions between epitopes and the CDR3β region of TCRs. Before making predictions for a specific pair of CDR3β and antigenic epitopes, it is essential to convert their alphabetical sequence information into numerical representations. In this research, the Atchley factors ^27^ were utilized as a method of numerical embedding for this purpose. The Atchley factors comprise five factors that encompass diverse physicochemical characteristics, enabling each amino acid (AA) to be effectively represented by these five factors. For each CDR3β-epitope pair, VitTCR translates it into a 3-dimensional numerical tensor referred to as AtchleyMap (**Methods**). As illustrated in **Figure 1a**, AtchleyMap has a width of 12 AAs, a height of 20 AAs, and a channel of 5, capturing the relevant information from the CDR3β and epitope sequences. The selection of the specific lengths (20 AAs for CDR3β sequences and 12 AAs for epitopes) is based on the results of statistical analysis (**Supplementary Figure 1**), which shows that approximately 98.89% of CDR3β sequences fall within the length range of 10 to 20 AAs, and 97.97% of epitopes fall within the length range of 8 to 12 AAs. Therefore, setting the lengths to 20 AAs and 12 AAs ensures that the majority of sequences are properly represented. To provide a more detailed description of the interactions, we partitioned the AtchleyMaps into distinct patches and assigned numerical labels to each patch accordingly (**Figure 1b, Methods**). As depicted in **Figure 1c**, VitTCR takes AtchleyMaps as inputs and generates predicted probabilities for “binding” or “no binding”, with the two predicted probabilities summing to 1. Notably, the patches numbered in **Figure 1b** correspond to the patches numbered in **Figure 1c** to ensure consistent interpretation.

**Figure 1.**
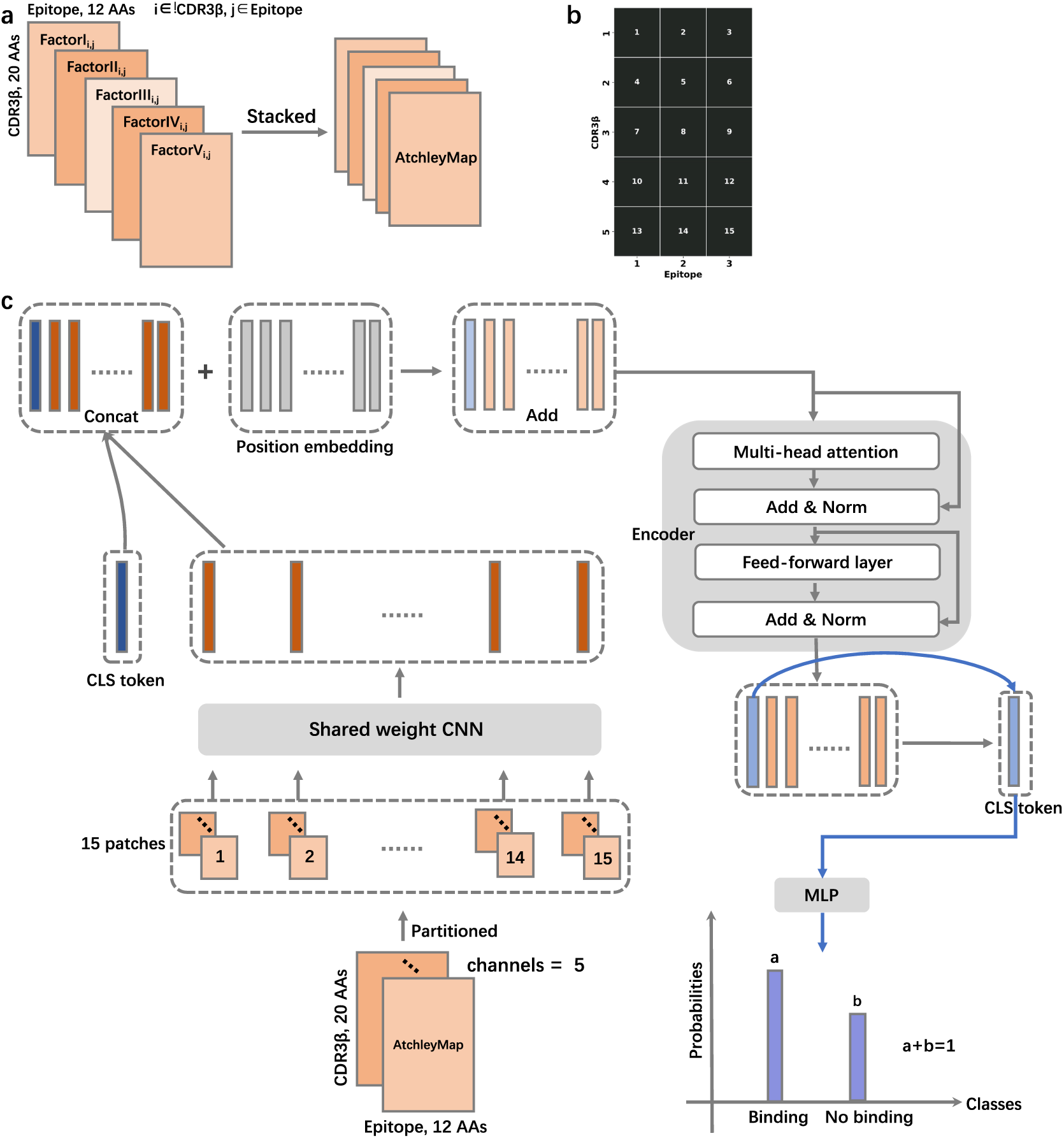
| The architecture of VitTCR for predicting interactions between CDR3βs and epitopes. **a**. Schematic diagram of AtchleyMap encoding. The value of each position was the absolute value of the difference in the Atchley factor of two corresponding AAs. The subscript i (ranging from 1 to 20) represents the position of amino acids of CDR3βs, and j (ranging from 1 to 12) represents the position of amino acids of epitopes. **b**. Strategy of patch division. Each AtchleyMap is partitioned into 15 patches. **c**. The architecture of VitTCR. VitTCR takes AtchleyMaps as inputs and generates predicted probabilities for “binding” or “no binding”, with the two predicted probabilities summing to 1. AA: amino acid; CNN: convolutional neural networks; CLS: classification; MLP: multilayer perceptron.

### Performance comparison with other methods

Following model optimization, we then conducted a comparative analysis between VitTCR and other published methods. The published methods for comparison involved NetTCR-1.0, NetTCR-2.0, ERGO_LSTM, ERGO_AE, and DlpTCR. These methods represent different architectures and techniques used for predicting interactions between CDR3βs and epitopes. Both NetTCR-1.0 and NetTCR-2.0 are traditional convolutional neural networks (CNNs), and they utilize convolutional layers to capture local patterns and features from input sequences. ERGO_LSTM employs a traditional long short-term memory (LSTM) network structure. LSTMs are recurrent neural networks (RNNs) that can capture long-term dependencies in sequential data. ERGO_AE utilizes an autoencoder, a type of neural network used for unsupervised learning. Autoencoders aim to reconstruct their input data, which helps in learning compressed and meaningful representations. DlpTCR is a model that integrates three different deep learning networks. It combines a CNN, an LSTM, and a fully connected neural network to capture both local and global features from the input sequences.

To ensure a fair comparison process, we employed the same training and test sets (**Supplementary Figure 2**), performed a fivefold cross-validation with five iterations for each method, and evaluated the classification performance of the trained models on the same independent test sets. As displayed in **Figure 2**, VitTCR outperforms these methods in terms of area under the receiver operating characteristic (AUROC), which provides strong evidence of the enhanced predictive power of VitTCR in accurately predicting interactions between epitopes and the CDR3β region of TCRs. The significant performance improvement observed in comparison to the existing methods underscores the advancements of VitTCR.

**Figure 2.**
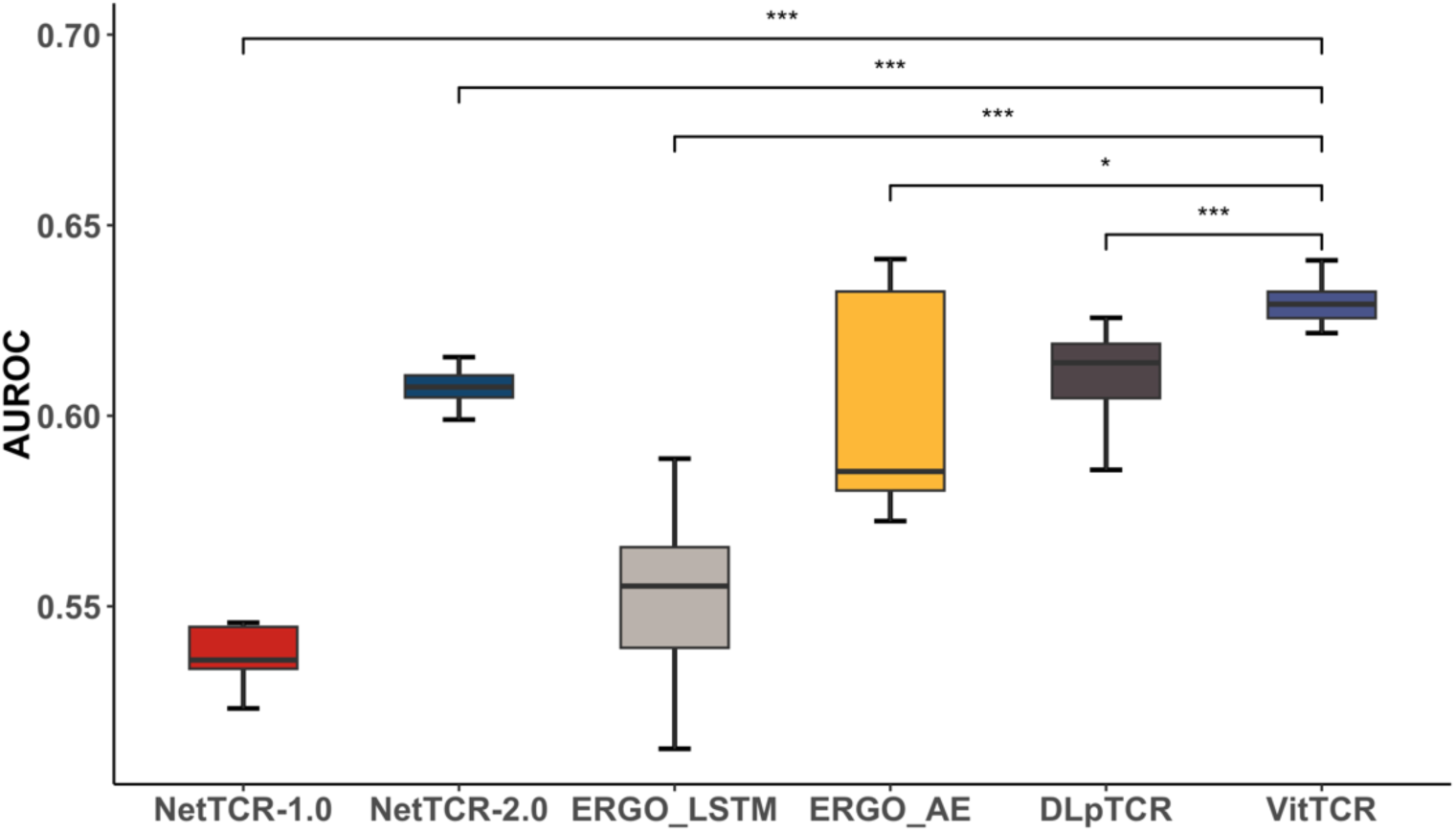
| Comparison of VitTCR with other published methods on an independent test set. All models were trained and validated using the same datasets. For each method, five repeated fivefold cross validations were conducted. These methods were compared in terms of AUROC.

### The influence of PBWM on model performance

To investigate positional bias and identify the crucial regions involved in the interactions between CDR3βs and epitopes, a set of 83 structurally resolved pMHC-TCR complexes was downloaded from the PDB database. The interacting AA pairs in these complexes were labelled using PyMOL (**Figure 3a, Supplementary Table 2**). The next step involved counting the occurrences of interacting AA pairs within the downloaded complexes and normalizing the counts for each patch (**Figure 1b**). This normalization was achieved by dividing the counts by the total number of interacting AA pairs. This analysis provided a positional bias weight matrix (PBWM). Each value in the weight matrix represents the percentage of the total number of interacting AA pairs occurring in that particular patch, and a higher value in a specific patch indicated a greater likelihood of CDR3β-epitope interactions occurring in that particular region. As displayed in **Figure 3b**, the sum of the percentages across the 15 patches equals 1, and most interactions tended to occur between the middle patch of the epitopes and the second to fourth patches of the CDR3β regions. In other words, the fifth to sixteenth AAs of CDR3βs and the fifth to eighth AAs of peptides contributed significantly to the interactions.

**Figure 3.**
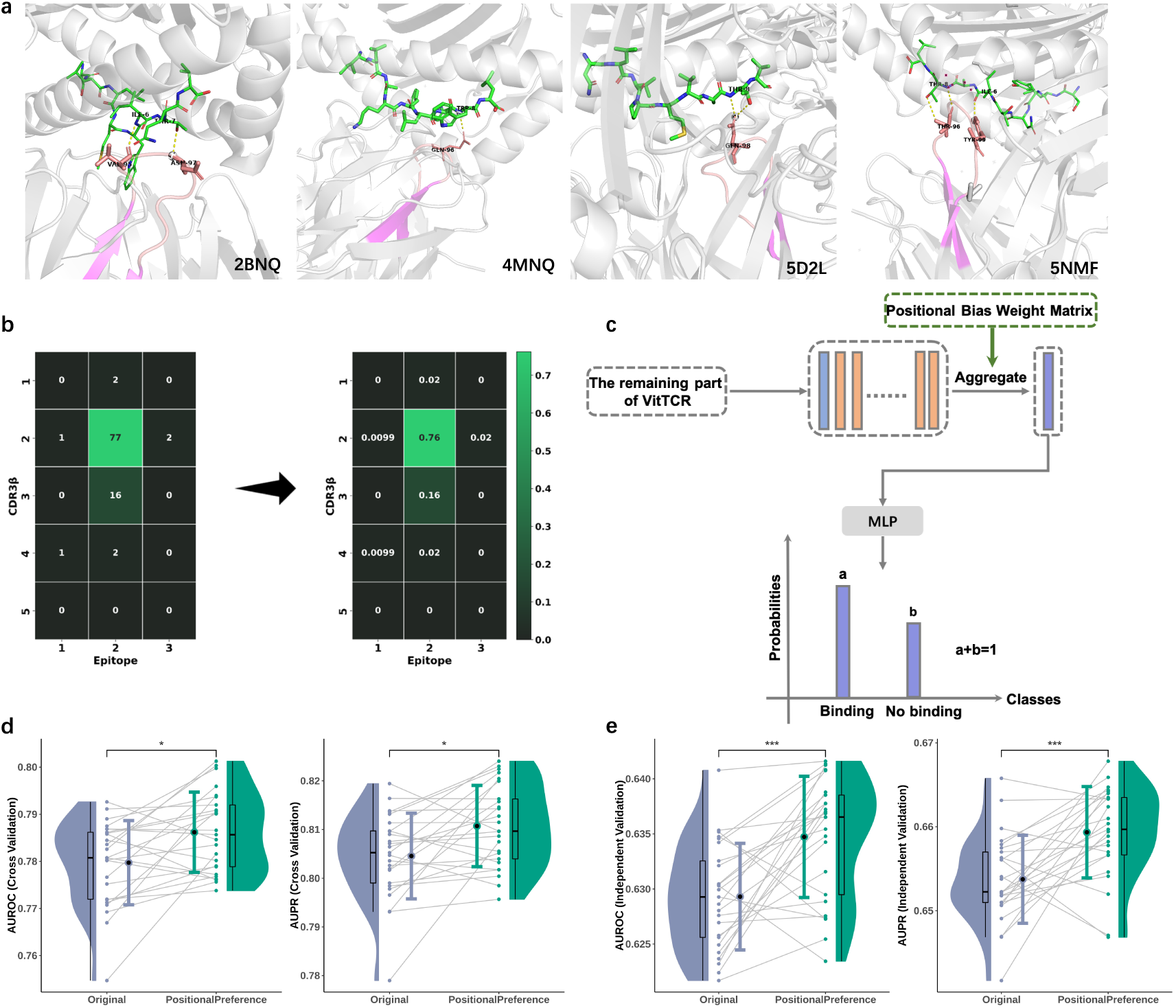
| The effects of PBWM on the performance of VitTCR. **a**. Four pMHC–TCR complexes (2BNQ, 4MNQ, 5D2L, and 5NMF) are labelled and coloured using PyMOL. The complexes are visualized with specific colour schemes: yellow dotted lines represent polar interactions in CDR3β-peptide pairs, the CDR3 region of TCRβ chains is coloured pink, and the epitopes are coloured green. **b**. The left panel illustrates the count of interacting AA pairs, where one AA is from the CDR3β region of the TCR and the other is from the epitope. A total of 101 AA pairs with interactions were identified. In the right panel, the matrix from the left panel is normalized by dividing the count in each patch by the total number of interacting AA pairs. **c**. Integration of PBWM with VitTCR. **d**. Comparison of the performance of VitTCR in the validation set before and after the integration of PBWM. **e**. Comparison of the performance of VitTCR in the independent test set before and after integration of the PBWM.

VitTCR provides a way of integrating PBWM (**Figure 3c, Methods**). To determine whether the PBWM can improve the prediction, we added the weight matrix into VitTCR and then compared the results of VitTCR with or without the PBWM. As illustrated in **Figure 3d** and **Figure 3e**, the performance of VitTCR was significantly improved on the validating set and testing set after adding the calculated weights, suggesting that the weight matrix captured the pattern of CDR3β-peptide interactions to some extent. However, we note that the sample size involved in calculating the PBWM was limited and insufficient to cover all cases. As more data become available, there is potential for further refinement and improvement of PBWM. This may contribute to the prediction of interactions between CDR3βs and epitopes.

### Cluster-based filtering can decrease false positives

For applications involving extensive experimental validation, it is essential to avoid false positive (FP) predictions since FP predictions will lead to unnecessary downstream validation and increased labour and time costs. Model sensitivity is usually not the main concern in practical applications for identifying TCR-peptide interactions. Instead, our primary goal is to increase the positive predictive value (PPV) of the model. The PPV is the proportion of true-positive (TP) results to the total number of predicted positive results (TP+FP). Thus, optimizing a model’s PPV becomes critical. Dash *et al*.^16^ found that CDR3βs with higher sequence similarities tend to recognize the same epitope. Therefore, we speculated that removing unclustered CDR3βs with distinct AA sequences from all other CDR3βs from the training dataset or testing dataset could improve the model performance. To test this hypothesis, we utilized iSMART^28^ for CDR3β clustering and removed unclustered CDR3βs from the dataset (**Methods**).

To investigate the effect of cluster-based filtering, we performed filtering operations on the training and test sets separately (**Supplementary Table 1**). First, we classified the trained models into four major categories, including Original (neither the training set nor the test set was filtered), Trainset-only (only training set clustered and filtered), Testset-only (only test set clustered and filtered), and Both-clustered (both training set and test set clustered and filtered). Then, we conducted five repeated fivefold cross validations under the four settings, and the model comparison results are shown in **Figure 4**. These results suggest that performing cluster-based filtering either on the training set or on the test set significantly improved the PPV of the model, and the best PPV was obtained for the category named Clustered. In addition, to ensure that this finding is not a coincidence, we also conducted the same analysis on other models, including NetTCR-2.0 and ERGO_AE. The results demonstrate that the application of cluster-based filtering to the dataset can significantly enhance the PPV, not only in our model but also in NetTCR-2.0 and ERGO_AE. Similar trends also occurred in terms of AUROC and AUPR (**Supplementary Figure 4**). These findings highlight the potential of cluster-based filtering as a valuable tool for optimizing the performance of prediction models. Additionally, we also conducted a downsampling evaluation and found that cluster-based filtering can reduce the dependence of the model on the data size (**Supplementary Figure 5**).

**Figure 4.**
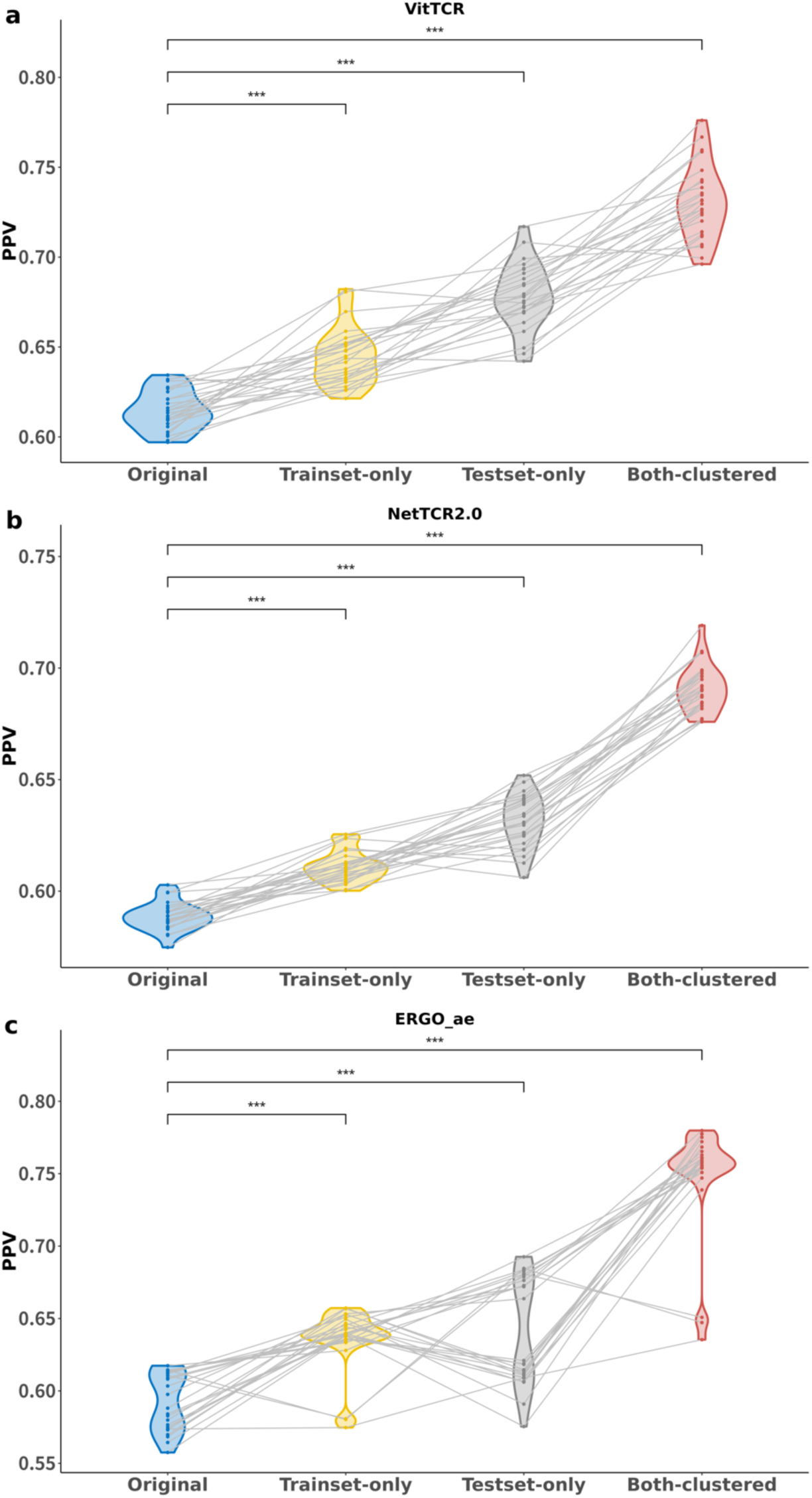
| The influence of cluster-based filtering on model performance in terms of PPV. According to the dataset configuration, the model was divided into four main categories: Original, Trainset-only, Testset-only, and Both-clustered. The PPVs of the three models were compared across the four different dataset settings. Five repeated fivefold cross validations were conducted under four settings, and a dot in this figure represents a fold replicate. **a**. Performance of VitTCR. **b**. Performance of NetTCR-2.0. **c**. Performance of ERGO_AE.

### Correlation between model predictions and TCR clone fraction

To verify model performance from a novel perspective, we conducted an investigation into the correlation between the model predictions and TCR clone fraction. Additionally, we conducted a comparative analysis of VitTCR with other methods, including ERGO_AE and NetTCR-2.0, using the same dataset. We obtained the single-cell TCR sequencing data of CD8^+^ T cells of four healthy human donors from the 10x Genomics platform ^29^. Given that the training dataset for our model was limited to HLA-A02:01 data points and HLA-A02:01 expression was observed only in Donor 1 and Donor 2, we focused the analysis on the data from these two donors. The dataset included a highly multiplexed panel consisting of 44 different pMHC multimers and 6 control pMHC multimers to determine the binding specificity of each CD8+ T cell. The affinity between each T-cell and each tested pMHC was quantified by counting the number of unique molecular identifier (UMI) sequences associated with that specific pMHC in the T cell. For each T cell, if the UMI count for any of the 44 pMHC multimers exceeded 10 and was more than five times higher than the highest UMI count for the 6 negative controls, the cell was considered to exhibit significant specificity towards that pMHC. During the cell filtration process, T cells showing significant specificity towards fewer than one pMHC or exceeding four pMHCs were excluded from the analysis. After the filtering of T cells, for each clonotype of TCR, we calculated the percentage of T cells harbouring that specific clonotype based on the total number of T cells measured in the donor, irrespective of the HLA-A*02:01 restriction. This percentage represents the clone fraction associated with that particular TCR clonotype and provides insights into the clonal expansion of T cells. Subsequently, for each TCR clonotype, the pMHC with the highest affinity was chosen, and the binding probability between the CDR3β of the TCR clonotype and the epitope of the pMHC was predicted. Finally, we calculated the Spearman correlation coefficient between clone fractions and predicted binding probabilities.

To ensure a rigorous and unbiased comparison process, we performed five repetitions of fivefold cross-validation for VitTCR, NetTCR-2.0 and ERGO_AE separately. Subsequently, we calculated the Spearman correlation coefficients between the predicted binding probabilities and the TCR clone fractions for each fold. This approach enabled us to objectively evaluate the associations between the predicted probabilities and the observed clone fraction for the methods mentioned above. As displayed in **Figure 6**, the predicted binding probabilities of all methods were significantly and positively correlated with the clone fraction. Higher predicted binding probabilities correspond to stronger clonal proliferation of T cells in the dataset. Each TCR clonotype was assigned 25 different predicted probabilities by VitTCR, NetTCR-2.0, and ERGO_AE. We calculated the mean predicted probability for each prediction method as the representative predicted probability for that particular TCR clonotype. **Supplementary Figure 6** displays the visualization of the mean predicted probabilities against the clone fractions. The relevant metrics are summarized in **Table 1**.

**Figure 5.**
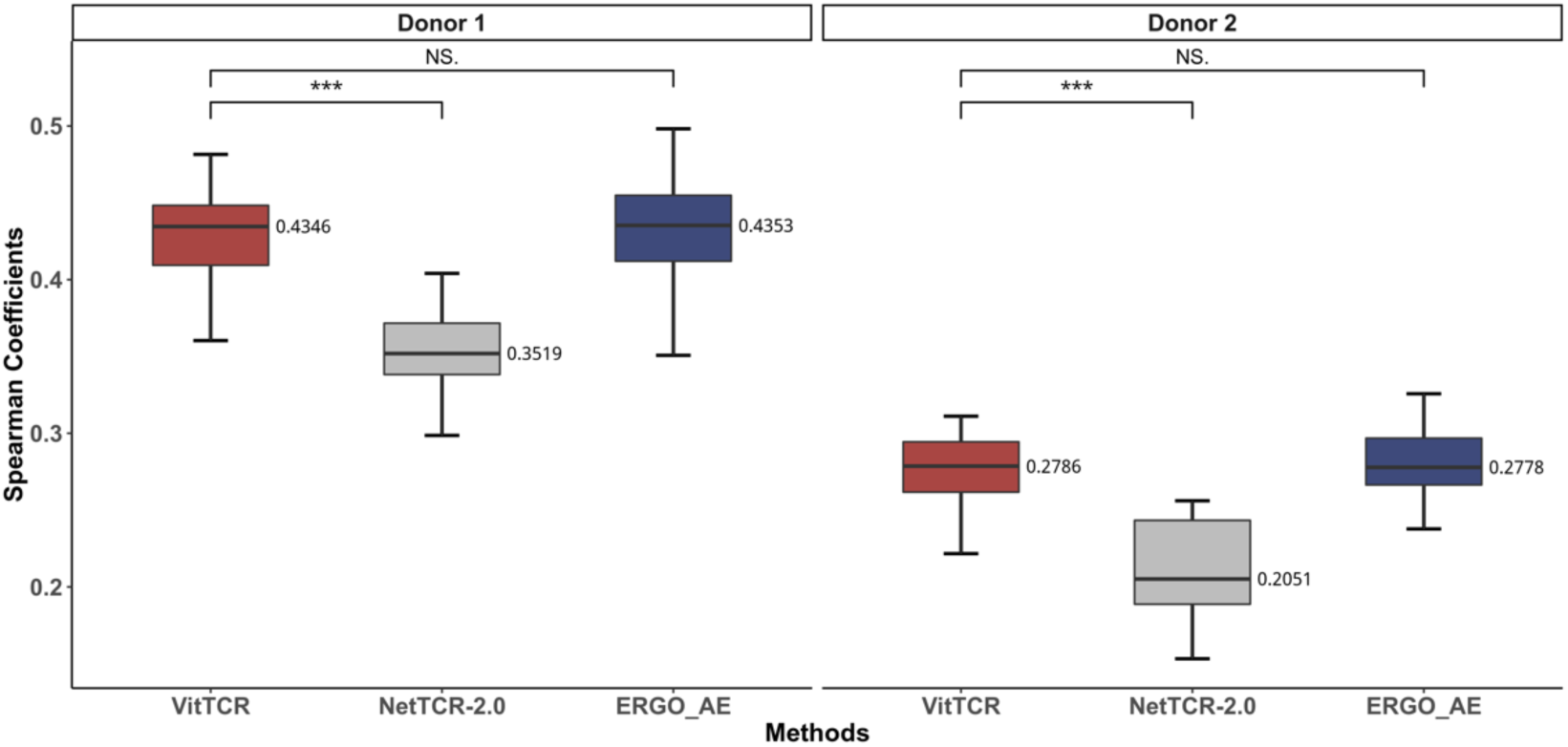
| Correlation between predicted probabilities and clone fractions for different methods. Spearman correlation coefficients between predicted probabilities and clone fractions were calculated for VitTCR, NetTCR-2.0, and ERGO_AE using different healthy donor data. For all methods, five repetitions of fivefold cross validation were performed. Each point of the boxplot represents the correlation coefficient between the predicted probabilities of the model and the clone fractions for each fold, and the number on the right side of the boxplot indicates the median correlation coefficient. The significance was determined using paired t tests.

**Figure 6.**
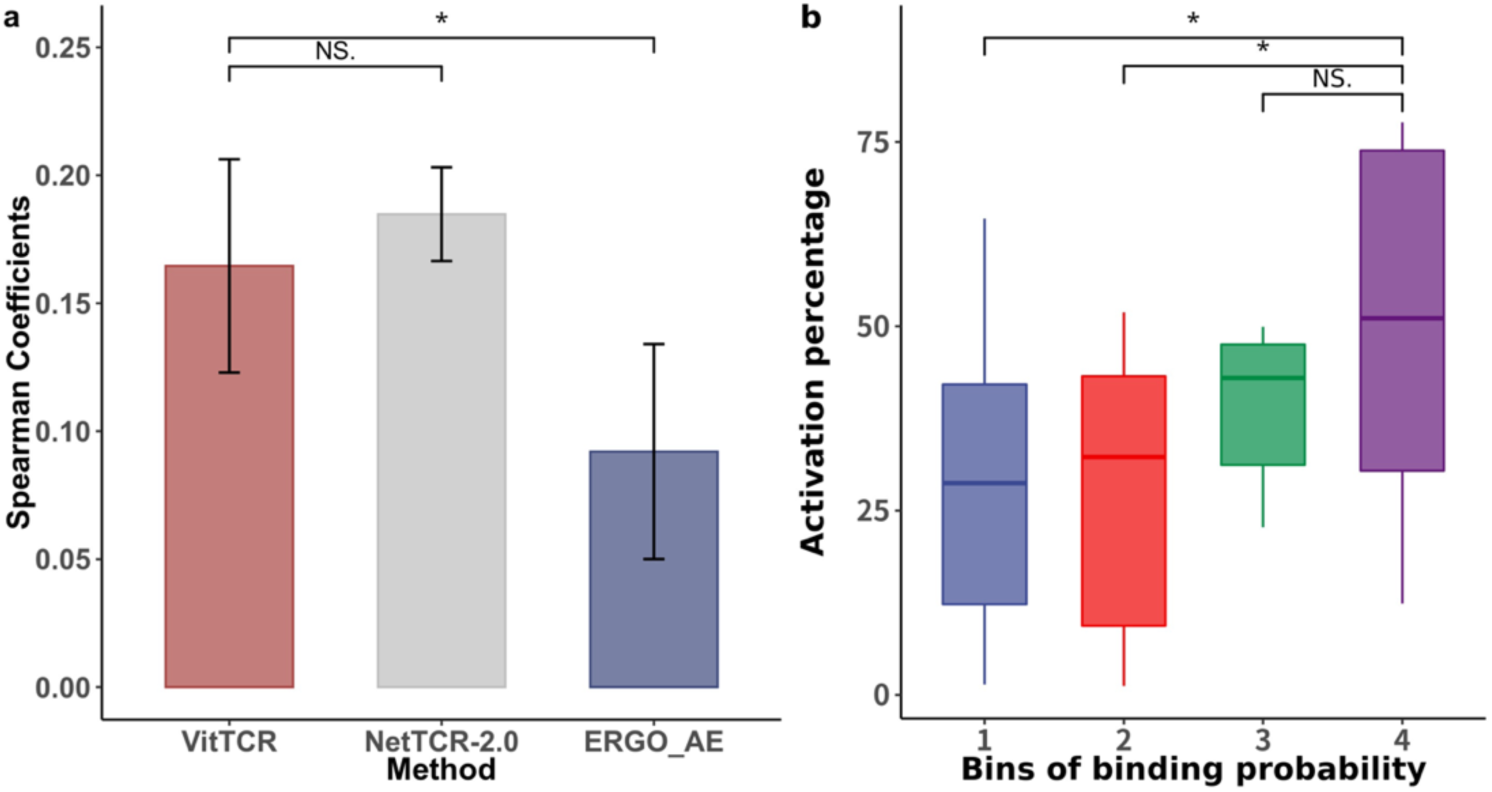
| Correlation between predicted probabilities and activation percentages for different methods. **a**. Spearman correlation coefficients between predicted probabilities and activation percentages of T cells were calculated for VitTCR, NetTCR-2.0, and ERGO_AE. For all methods, fivefold cross-validation was performed. Each point of the boxplot represents the correlation coefficient between the predicted probabilities of the model and the activation percentages for each fold. The significance was determined using paired t tests. **b**. The binding probabilities ranging from 0 to 1 were divided into five bins, with each bin representing an interval of 0.2. The x-axis represents the bins of binding probabilities predicted by VitTCR, while the y-axis represents the activation percentages of TCR clonotypes.

Specifically, the correlation coefficient between the predicted binding probabilities and the clone fraction was significantly higher for VitTCR than for NetTCR-2.0. In the comparison between VitTCR and ERGO_AE, VitTCR exhibited a higher median correlation coefficient. Generally, VitTCR, in particular, demonstrates superior performance compared to NetTCR-2.0 and ERGO_AE in reflecting clonal proliferation based on predicted binding probabilities.

### Correlation between model predictions and T-cell activation

Furthermore, we sought to examine whether the predicted binding probabilities were positively correlated with the activation percentage of T cells. Additionally, we conducted a comparative analysis of VitTCR with other methods, including ERGO_AE and NetTCR-2.0, using the same dataset. In a previous study (manuscript submitted), we cocultured 15 T-cell clonotypes with 11 immunogenic peptides of SARS-CoV-19 in a pairwise manner and selected CD137 as a marker of T-cell activation to quantify the percentage of T-cell activation. In total, we obtained 165 CDR3β-epitope pairs. The aforementioned steps of coculture and quantification of T-cell activation were repeated three times so that each CDR3β-epitope pair had three activation percentage values, and the mean value was taken as its percentage activation in this study.

As depicted in **Figure 6a**, the Spearman correlation coefficient between the predicted binding probabilities and the activation percentages of T cells was significantly higher for VitTCR than for ERGO_AE, while there was no significant difference in performance between VitTCR and NetTCR-2.0. As shown in **Figure 6b**, CDR3β-epitope pairs with higher T-cell activation percentages tended to have higher predicted binding probabilities by VitTCR. This observation suggests that the predicted results of VitTCR may partially reflect cell activation, indicating its potential role in capturing the activation status of T cells.

## Discussion

In this study, we developed a novel method named VitTCR (**Figure 1c**) to predict the interactions between epitopes and the CDR3β region of TCRs. For each CDR3β-epitope pair, VitTCR encodes it into a 3-dimensional numeric tensor named AtchleyMaps. The length of the tensor corresponds to the length of CDR3β (20 AAs), the width corresponds to the length of the epitope (12 AAs), and the channels represent the 5 Atchley factors. Subsequently, VitTCR takes the AtchleyMaps as inputs and outputs the predicted binding probability for each CDR3β– epitope pair. Compared with other published methods, VitTCR outperforms these methods in predicting interactions between epitopes and the CDR3β region of TCRs, suggesting the enhanced predictive power of our method (**Figure 2**).

The flexibility of the contact between the AAs of CDR3βs and epitopes is constrained by the antigen presentation of MHCs and the overarching spatial structure of the pMHC-TCR complex. Thus, the contact probabilities between the AAs of CDR3βs and epitopes are different due to their positions on the AA sequences. To investigate positional bias and identify the crucial regions involved in the interactions between CDR3βs and epitopes, we downloaded 83 pMHC-TCR complexes from the PDB database and counted the interacting AA pairs in each patch. Normalization was performed by dividing the counts of interacting AA pairs in each patch by the total number of interacting AA pairs. Afterwards, we obtained a matrix named PBWM. To make use of this information, VitTCR provides options for the integration of PBWM (**Figure 3c**). The performance of VitTCR was significantly improved on the validating set and independent test set after adding the PBWM, suggesting that the weight matrix captured the pattern of CDR3β-peptide interactions to some extent. Notably, the sample size involved in calculating the PBWM was limited and insufficient to cover all cases. As more data become available in the future, the PBWM can be further refined and enhanced.

We also found that performing cluster-based filtering on the training and/or test sets with iSMART can reduce noise in the data and improve the PPV (**Figure 4**). The improvement brought by cluster-based filtering was also seen in NetTCR-2.0 and ERGO_AE. Additionally, downsampling analyses suggest that cluster-based filtering can decrease the model’s dependence on the data size. The results suggest that cluster-based filtering on datasets has the potential to be a valuable tool for optimizing model performance.

To assess the generalizability of the prediction models, we also investigated the correlation between the model predictions and TCR clone fraction as well as T-cell activation percentage. Regarding the TCR clone fraction, we used the single-cell TCR sequencing data of CD8^+^ T cells acquired from two healthy human donors. The binding specificities of each CD8^+^ T cell were assessed by a highly multiplexed panel of distinct pMHC multimers.

Spearman correlation between the predicted binding probabilities and clonal fractions of TCR clonotypes suggests that, in general, the stronger the interactions between T cells and peptides are, the stronger the clonal expansion of T cells. Additionally, we sought to evaluate model performance by examining predicted probabilities in relation to the activation percentages of T cells. To do this, we employed a dataset comprising 15 TCR clonotypes and 11 peptides, which were identified from individuals who had recovered from SARS-CoV-19. T cells with a specific clonotype and peptide were cocultured pairwise, and the activation percentages of T cells were quantified by flow cytometry. The Spearman correlation coefficient between the binding probabilities and T-cell activation percentages suggested that the predicted results of VitTCR may partially reflect cell activation.

Generally, VitTCR is reliable for CDR3β-peptide interaction prediction. In addition, VitTCR provides an option for adding PBWM. As more data appear, the positional bias weights will be more detailed, thus increasing the generalizability of the model. In this study, due to the insufficient paired data of TCR, we only considered CDR3β without considering CDR3α. However, CDR3α also plays a role in peptide recognition. By considering CDR3α, predictions will be more precise and reliable. We expect more training data on TCR–pMHC, which will help us to make predictions more reliable and provide an effective adjunct for vaccine design and tumour immunotherapy.

## Methods

### Statistical analysis of sequence length

CDR3β plays a crucial role in the recognition of antigens by directly binding to antigenic epitopes. Consequently, when encoding the interactions between the TCR and antigenic epitopes, we focus solely on the CDR3β region of the TCR. Considering the specific attributes of VitTCR, it is essential to maintain a consistent input shape for the model. To determine the optimal length threshold, a thorough statistical analysis was performed on the lengths of CDR3β and antigenic epitopes within databases. As depicted in **Supplementary Figure 1**, a substantial majority, precisely 98.89%, of the entire set of CDR3β sequences exhibit lengths spanning from 10 to 20 amino acids. Likewise, a substantial proportion, amounting to 97.97%, of all epitopes exhibited lengths within the range of 8 to 12 amino acids. Therefore, for CDR3β sequences shorter than 20 amino acids or epitopes shorter than 12 amino acids, zero padding will be applied from the N-terminal to the C-terminal end of the sequences.

### Dataset processing

To train the model, we collected experimentally verified CDR3β-peptide pairs as positive samples from IEDB and McPAS. The data from VDJdb were utilized as a testing dataset to validate the model. All negative samples were generated by mismatching the peptide in each positive sample with a randomly selected CDR3β sequence from a healthy donor in TCRdb. Furthermore, we focused on the CDR3β sequences with 10-20 AAs, which started with ‘C’ AAs and ended with ‘F’ or ‘W’ AAs, and the peptides with 8-12 AAs that were presented by human MHC class I molecules. Therefore, we obtained an HLA-A*02:01-restricted training set with 38,712 data points before cluster-based filtering and 28,584 data points after cluster-based filtering using iSMART according to our experimental needs. The number of data points in the testing set was 5,250. Additionally, TCR sequences of CD8^+^ T cells acquired from healthy human donors (18,331 and 8,337 cells from Donor 1 and Donor 2, respectively) were used to measure the generalization of VitTCR by calculating correlations between the predicted binding probabilities and clonal fractions. To demonstrate the reliability of VitTCR, a total of 165 experimentally validated CDR3β-peptide pairs from COVID-19 recoverees were also used to display a positive correlation between the predicted binding probabilities and the activation percentages of T cells.

### Atchleymap encoding

Prior to predicting a pair of CDR3β and antigenic epitopes, it is necessary to convert their sequence information into numerical representations. In this study, we employed Atchley factors to encode CDR3β and antigenic epitopes. The Atchley factors consist of five factors that represent different physicochemical characteristics, with each amino acid being characterized by these five factors. For each factor, we encoded them using the principle shown in **Eq. 1**: the subscript ***i*** (ranging from 1 to 20) represents the position of amino acids of CDR3βs; ***j*** (ranging from 1 to 12) represents the position of amino acids of epitopes; and the coordinates [**i**, **j**] correspond to the absolute difference between the values of the Atchley factor of the two amino acid residues. Each Atchley factor generates a separate map, resulting in a total of five maps. These maps are then stacked together, and the resulting output tensor is referred to as AtchleyMap.

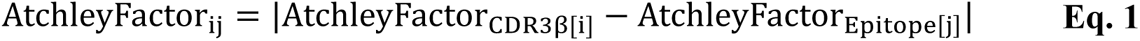

### Patch division

To provide a more detailed representation of the interactions, we partitioned the AtchleyMaps into distinct patches and assigned numerical labels to each patch accordingly (**Figure 1b**). Prior to the partitioning process, CDR3β sequences with fewer than 20 amino acids and epitopes with fewer than 12 amino acids were zero-padded from the N-terminus to the C-terminus. Consequently, we obtained 15 patches, each with a side length of 4 amino acids.

### Architecture of VitTCR

As depicted in **Figure 1c**, VitTCR took the encoded AtchleyMaps as input and generated predicted probabilities as output. Initially, VitTCR applied a two-dimensional convolutional layer with 256 kernels (size of 4 × 4 and stride of 4) to extract informative features from AtchleyMaps. This process can also be seen as dividing the encoded AtchleyMaps into 15 patches, where each patch was vectorized into a vector of size [256] using the two-dimensional convolutional layer (size of 4 × 4 and stride of 4). After the convolution step, these vectors were concatenated into a two-dimensional vector of size [15, 256]. Subsequently, the two-dimensional vector was concatenated with a randomly generated one-dimensional vector (size of [1, 256]), which serves as a learnable classification (CLS) token. At this point, the concatenated vector, which contained 16 tokens, had a size of [16, 256]. To retain the relative positional information, a position embedding of the same size as the concatenated vector was added to the vector. Afterwards, an encoder that consisted of a normalization layer, a multi-head attention layer, a dropout layer, and a feedforwards layer was used to further encode the 16 tokens. Finally, a multilayer perceptron (MLP) layer with two units was added on top of the encoder to extract features from the CLS token, and a softmax function was applied in the final output layer to export a vector P of size [2, 1]. The shape of P is the number of classes, including “binding” and “no binding”, with each element representing the probabilities related to the class labels, and the predicted probabilities summed to 1.

### Self-attention mechanism of VitTCR

The self-attention mechanism plays a crucial role in enabling VitTCR to predict the specific recognition between a CDR3β-epitope pair. As depicted in **Figure 1**, the Encoder module in VitTCR is similar to the Encoder module in Transformer ^30^. In the encoder module, the process of encoding a specific token, denoted as Token_i (where i represents the token number, i ∈ [CLS,1,2…,15]), primarily consists of three parts. The first part involves creating three vectors based on the input Token_i: a query vector (q_Token_i_), a key vector (k_Token_i_), and a value vector (v_Token_i_). These vectors are obtained by multiplying the vectorized tokens with the query matrix (W^Q^) as described in **Eq. 2**, the key matrix (W^K^) as described in **Eq. 3**, and the value matrix (W^V^) as described in **Eq. 4**. In VitTCR, to generate the three matrices mentioned above, a fully connected layer is initialized for all input tokens [N, 256], thereby increasing the number of channels in the resulting output to three times the original number [N, 768]. Here, N represents the total number of tokens, which is 16 for VitTCR. Subsequently, the output is reshaped to [3, num_heads, N, C], where num_heads denotes the number of heads of the multihead attention (set to 4 in VitTCR), and C represents the number of channels (calculated as the quotient of 256 divided by num_heads, yielding 64). Finally, the reshaped output [3, num_heads, N, C] is divided into three parts along the first dimension, resulting in W^Q^, W^K^, and W^V^, with all three matrices having the shapes of [num_heads, N, C]. In the second part, Score_Token_i_ is calculated. Score_Token_i_ is a list of length 16, where each element determines the level of attention that should be assigned to the other tokens during the encoding process of Token_i. As described in **Eq. 5**, Score_Token_i_ is obtained by computing the dot products between the query vector q_Token_i_ of Token_i and the key vectors k_Token_j_ of the other tokens Token_j (where j ∈ [CLS,1,2,…,15]). To prevent model collapse, these dot products are divided by the square root of the number of channels in W^Q^ (since W^Q^, W^K^, and W^V^ have an equal number of channels, which is 64). Finally, a softmax normalization operation, as shown in **Eq. 5**, is applied to these values. This normalization ensures that the sum of all the normalized values is equal to 1, as depicted in **Eq. 6**. The third part of the encoding process is to calculate the weighted average of the value vectors v_Token_j_ for all tokens based on their corresponding scores Score_Token_i_[j], as shown in **Eq. 7**. The resulting weighted average, denoted as Embedded_Token_i_, represents the encoded representation of the token Token_i based on the self-attention mechanism.

The functioning of the self-attention mechanism is elucidated in **Supplementary Figure 3**. Taking the encoding process of a CLS token as an illustrative example, the scores between the CLS token and the other tokens are visually represented by the thickness of the connecting lines. A thicker line signifies a higher score, while a thinner line indicates a lower score. The objective is to assign higher scores and greater weight to tokens that carry relevant information while assigning lower scores and lesser weight to tokens that contain irrelevant noise. This step is instrumental in preserving meaningful information while disregarding irrelevant noise. Although the SoftMax score between a token and itself tends to be the highest, it remains important to consider other tokens that are interconnected with the current token to capture their contextual relevance.

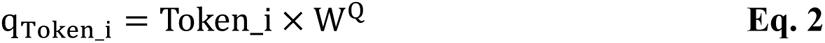

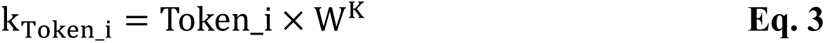

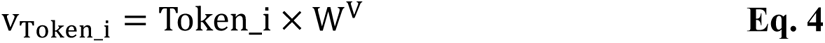

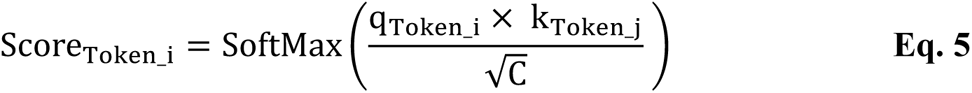

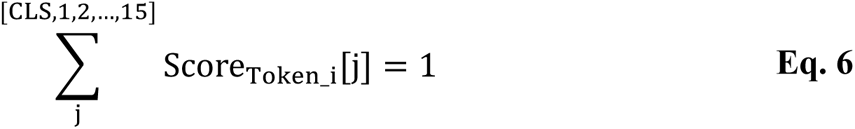

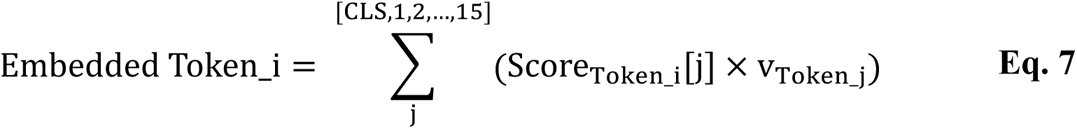

### PBWM integration

In VitTCR, the integration of PBWM involves the calculation of the mean value between Score_Token_CLS_ and PBWM (**Eq. 8**), followed by the calculation of the embedded CLS token, as illustrated in **Eq. 9**. To evaluate the influence of incorporating PBWM on the predictive accuracy of the model, a comparative analysis was conducted between VitTCR with the inclusion of the computed PBWM and VitTCR without the incorporation of this additional weight matrix.

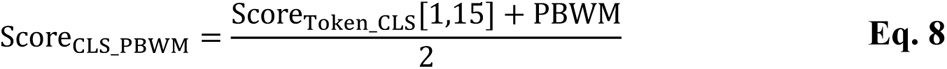

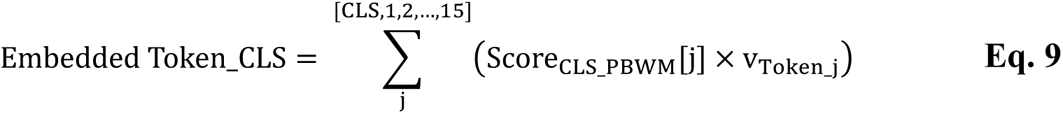

### Cluster-based filtering

iSMART was utilized for cluster-based filtering of CDR3βs. The command line for iSMART is “**python iSMARTv3.py-f cdr3β_total.txt-v**”. During the clustering process of N CDR3β sequences, iSMART calculated the pairwise alignment score based on the BLOSUM62 matrix and normalized the score by the length of the longer CDR3β sequence. Then, an N-by-N pairwise scoring matrix was obtained. Finally, the cut-off value was set as 3.5 to filter out all the unclustered CDR3βs.

## Supplementary Figure

**Supplementary Figure 1.**
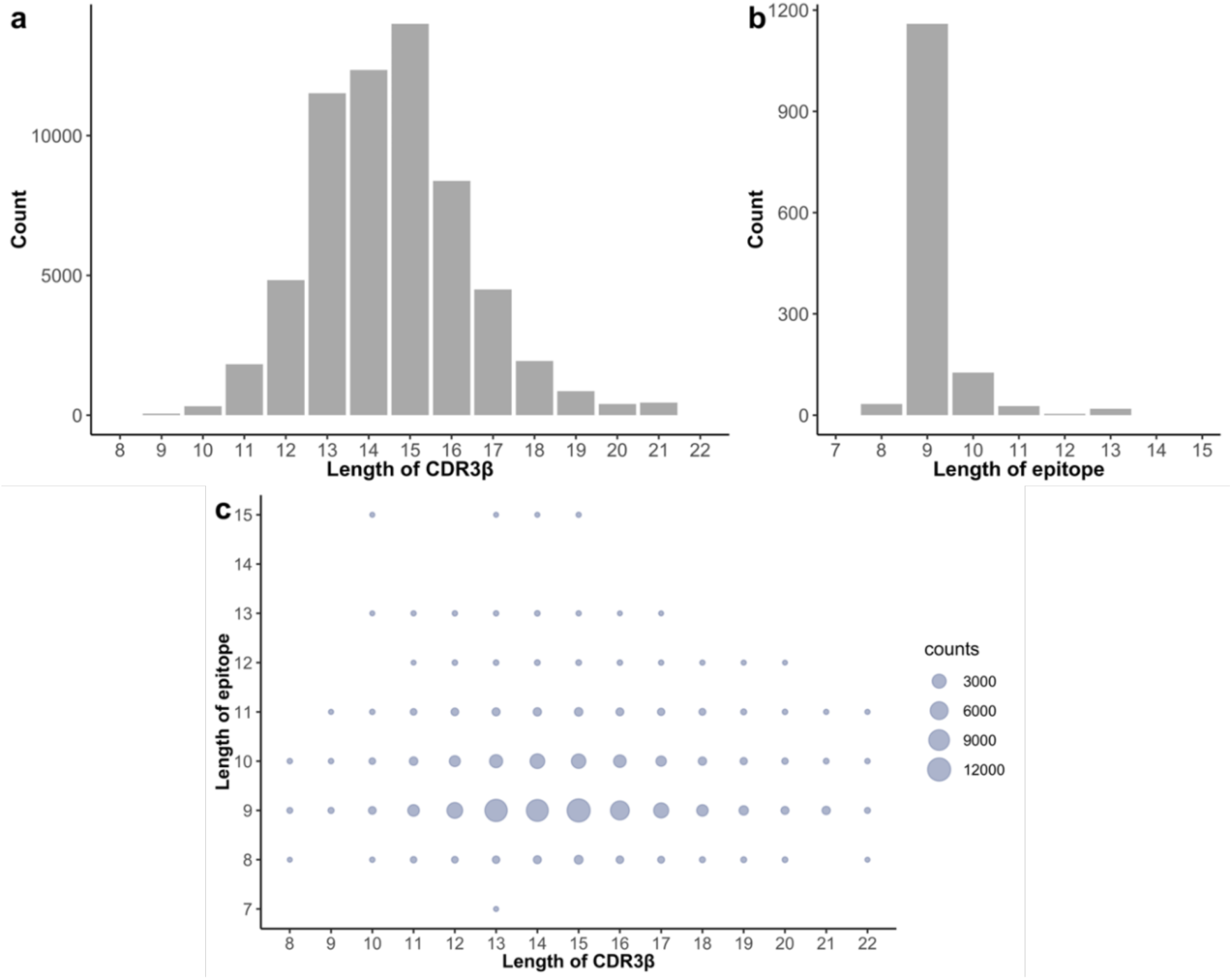
| Statistics on the lengths of CDR3β and epitopes in existing databases. **a.** The distribution of sequence lengths of CDR3β shows that the majority of CDR3β sequences consist of 10 to 20 amino acid residues. **b.** The distribution of sequence lengths of epitopes reveals that most antigenic epitopes comprise 8 to 12 amino acid residues. **c.** The distribution of sequence length combinations of CDR3β-epitope pairs in public databases is depicted in the graph. Each dot on the graph represents a specific sequence length combination, where the x-axis corresponds to the sequence length of CDR3β in a CDR3β-epitope pair, and the y-axis corresponds to the sequence length of the epitope in the same pair. Notably, the combination of 15 amino acid residues for CDR3β and 9 amino acid residues for the epitope exhibited the highest number of samples.

**Supplementary Figure 2.**
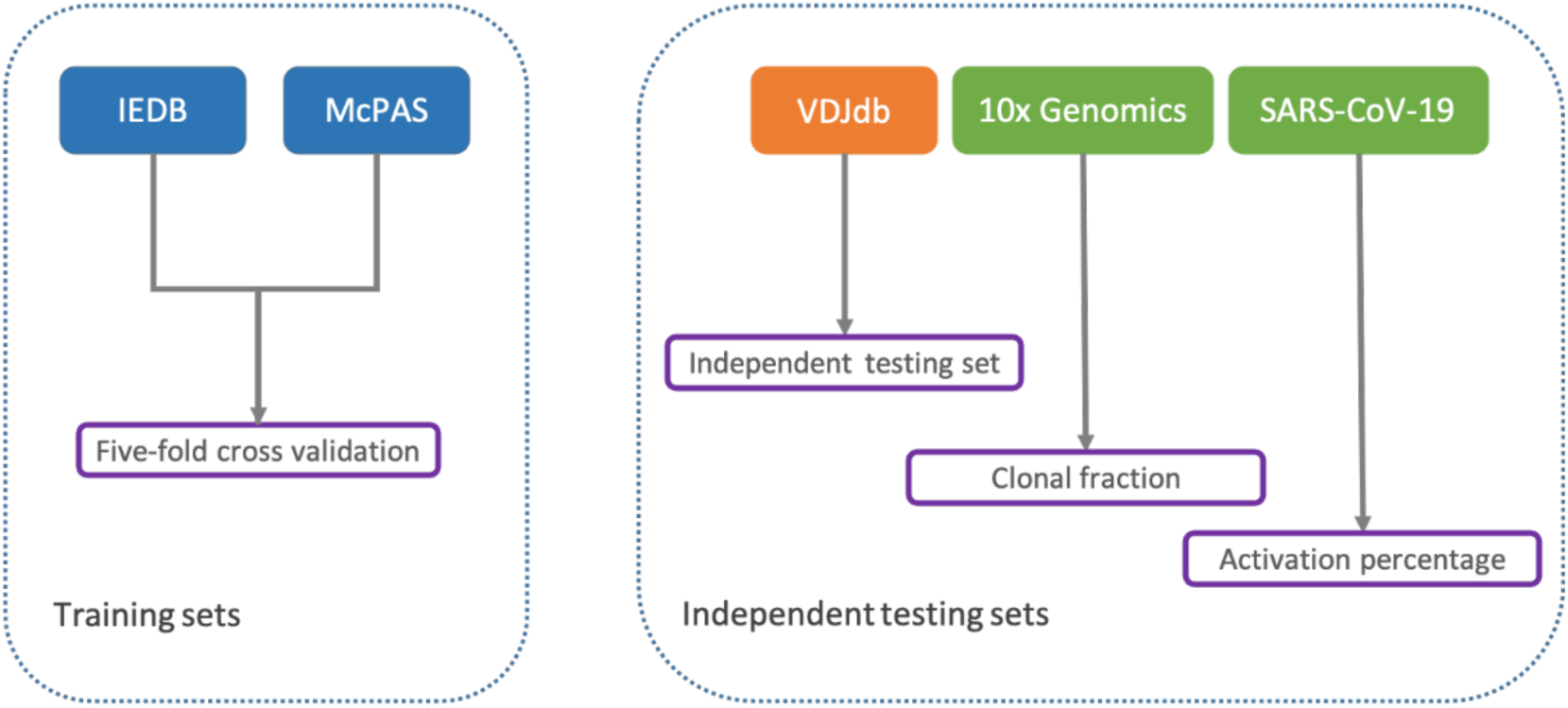
| The schematic diagram illustrates the datasets utilized for model training. Samples from IEDB and McPAS were utilized as the training set, while samples from VDJdb served as an independent test set. The model was further validated based on clonal fraction using samples from 10x Genomics and activation percentage using samples from SARS-CoV-2.

**Supplementary Figure 3.**
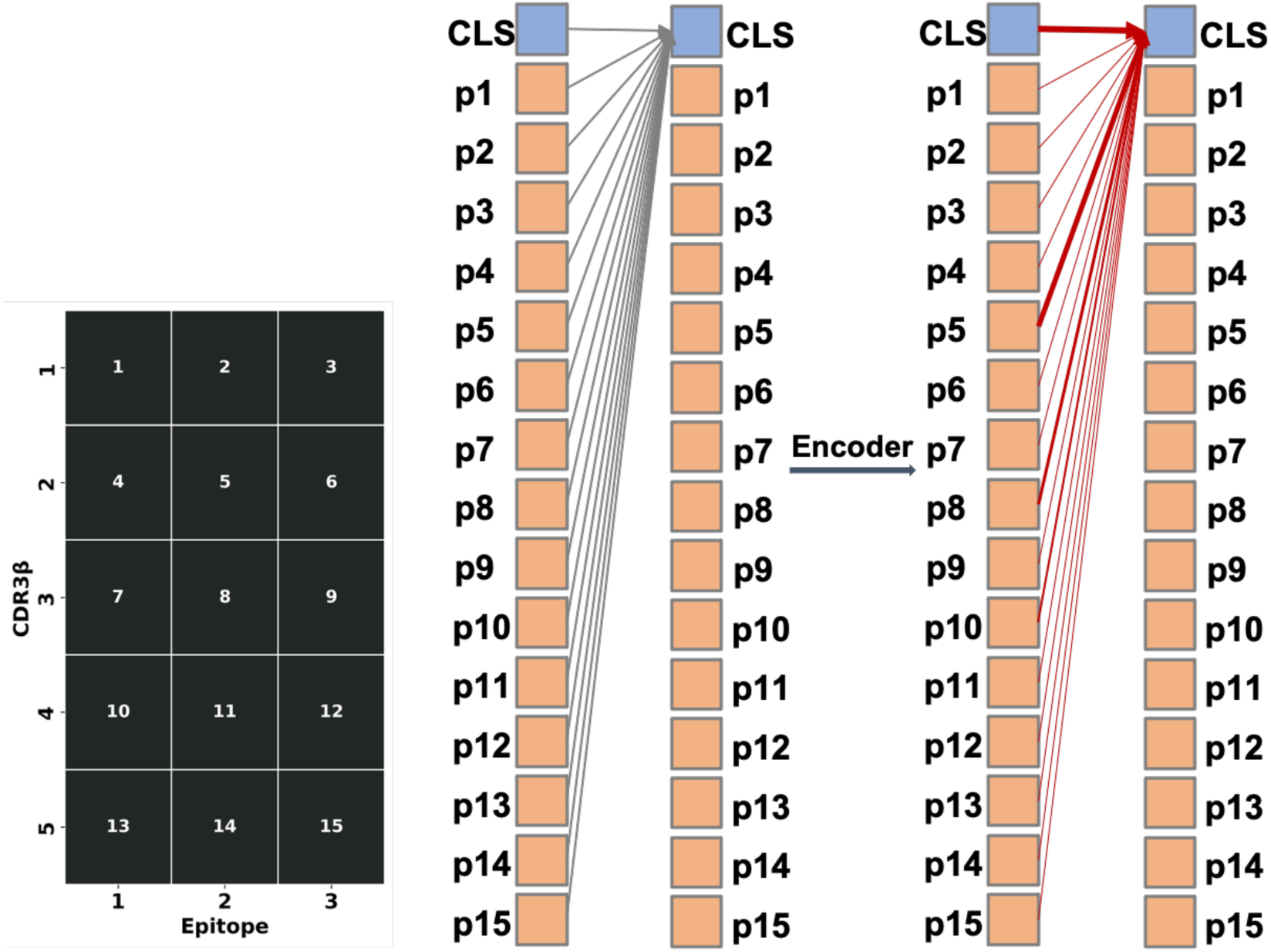
| Illustration of how the self-attention mechanism influences prediction. The patches numbered in the left panel correspond to the tokens numbered in the right panel. In the right panel, the lines connecting any two tokens represent correlation. Before the Encoder module, the scoring between CLS tokens and other tokens exhibits randomness (indicated by grey lines), while after the Encoder module, the scoring between CLS tokens and other tokens becomes more varied, with some scores being high (represented by thick red lines) and others being low (represented by thin red lines). The thickness of the lines reflects the strength of the score between the two tokens, where thicker lines signify higher scores and thinner lines represent lower scores.

**Supplementary Figure 4.**
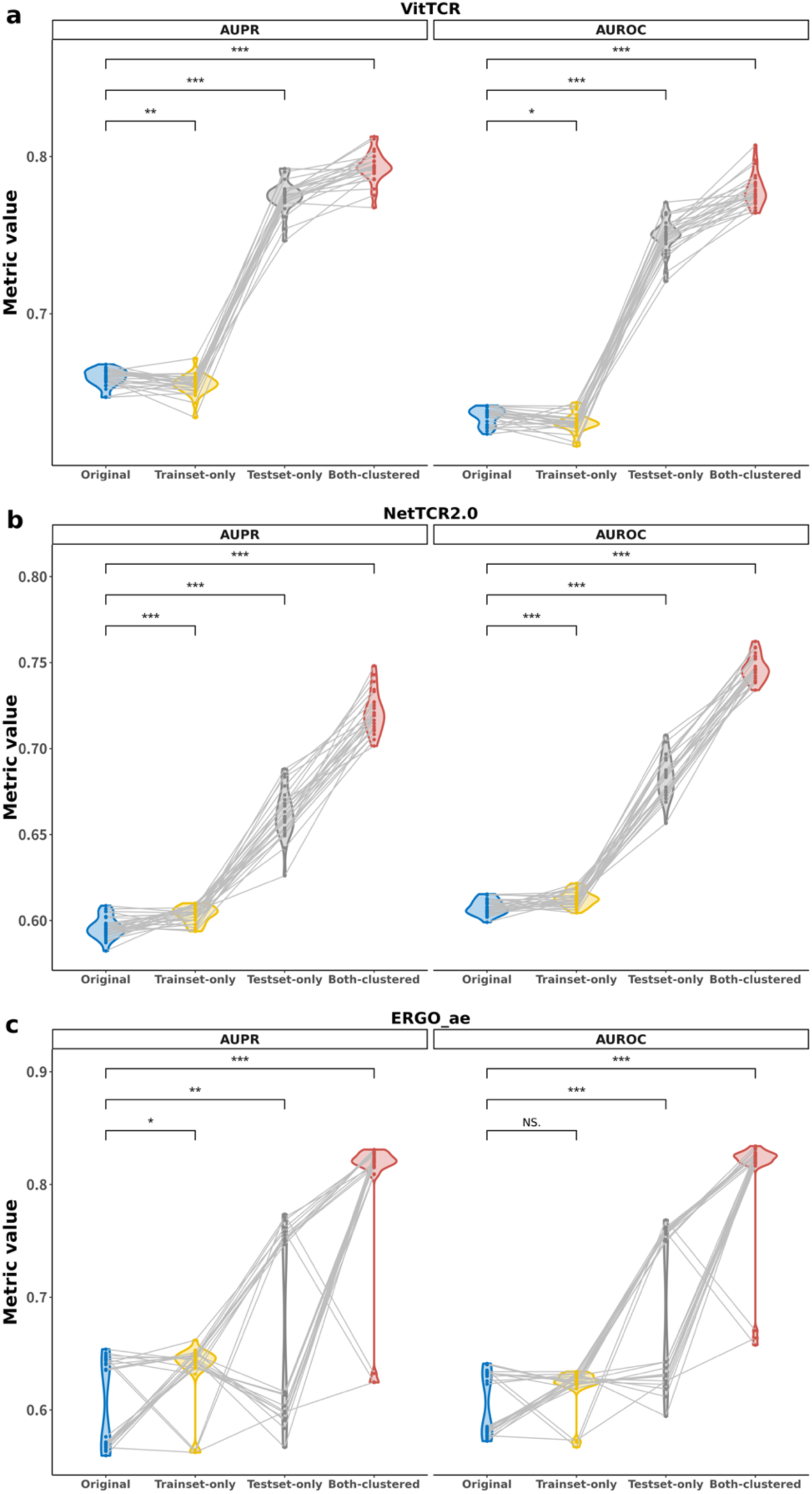
| The influence of cluster-based filtering on model performance in terms of AUROC and AUPR. The AUROC and AUPR of the three models were compared across the four different dataset settings. Five repeated fivefold cross validations were conducted under four settings, and a dot in this figure represents a fold replicate. **a**. Performance of VitTCR. **b**. Performance of NetTCR-2.0. **c**. Performance of ERGO_AE.

**Supplementary Figure 5.**
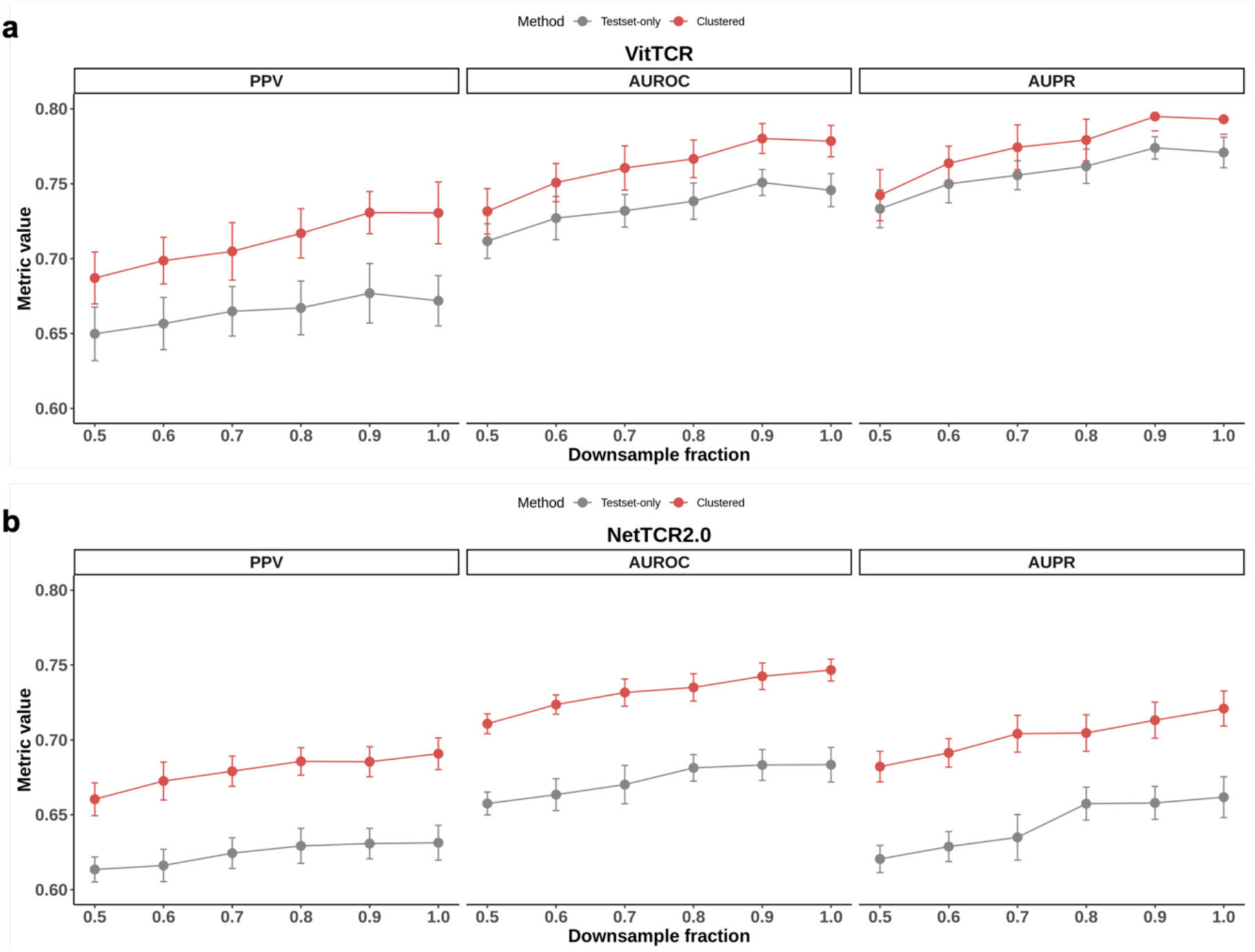
| The impact of cluster-based filtering on model performance in relation to its dependence on the data size. We performed downsampling on the training set, varying the downsampling ratios from 0.5 to 0.9 in increments of 0.1. For clarity, we focused on two specific settings: “Testset-only” and “Clustered”. The two settings utilized the same independent test set while employing different training sets, one with clustering-based filtering and the other without. The downsampling analysis indicates that cluster-based filtering has the potential to decrease the reliance of the model on the size of the dataset.

**Supplementary Figure 6.**
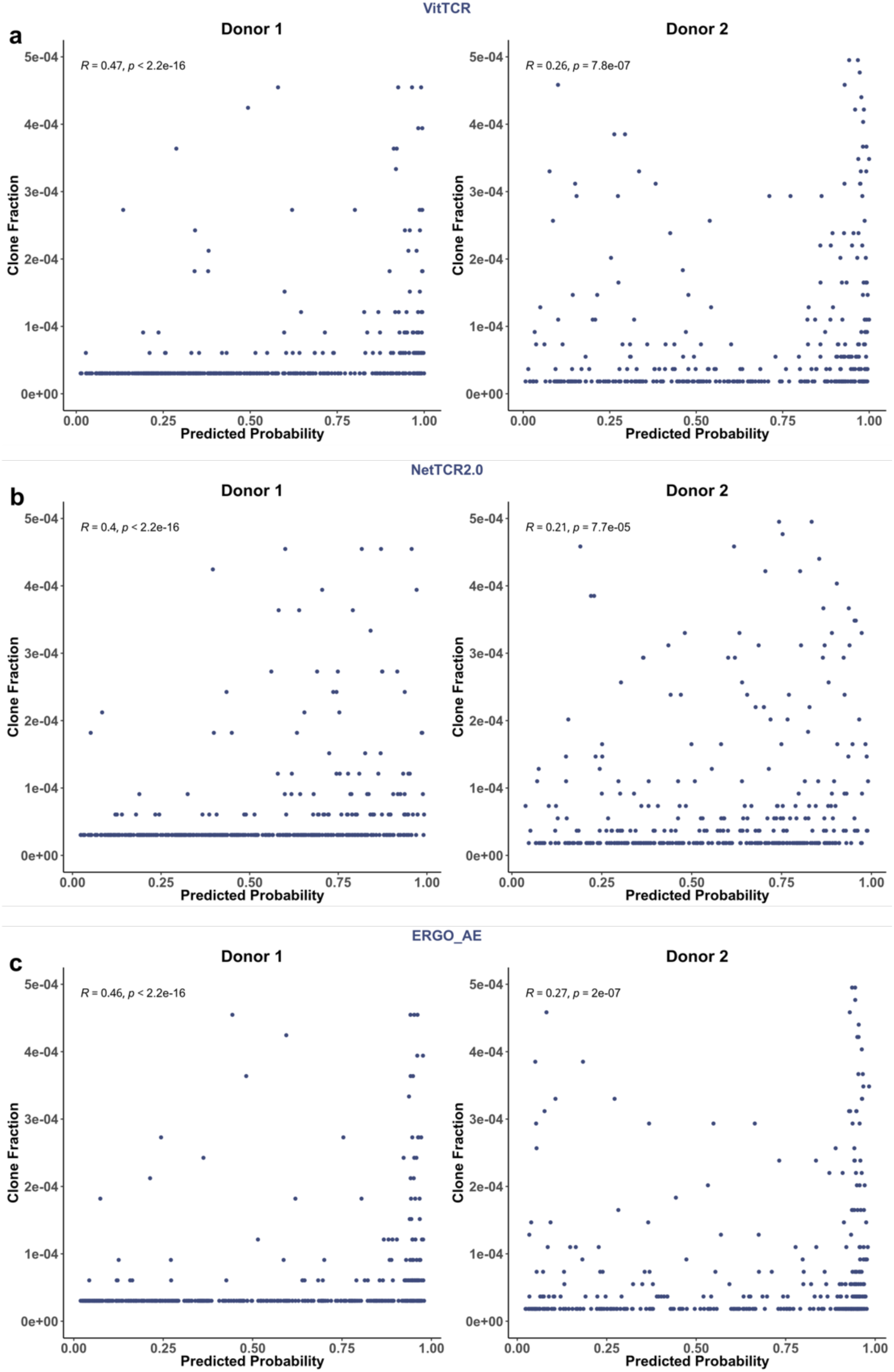
| Influences of cluster-based filtering on model performances. A dot in the scatter plot represents a TCR clonotype. The x-axis representing the mean of predicted probabilities and the y-axis representing the clone fraction. The Spearman correlation coefficient (R) and significance (p) are also indicated in figure. **a**. Performance of VitTCR. **b**. Performance of NetTCR-2.0. **c**. Performance of ERGO_AE.

