## Supplementary Figures and Tables for "VitTCR: A deep learning method for peptide recognition prediction"

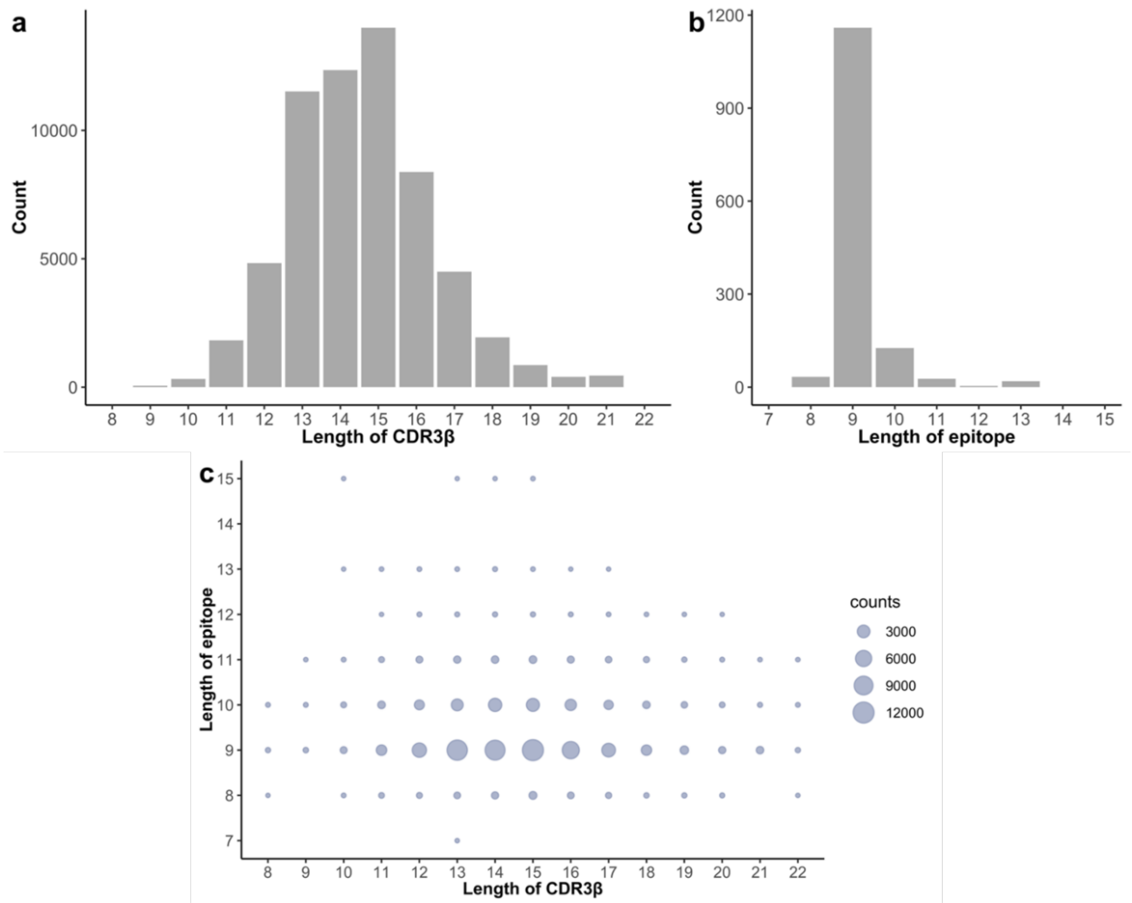


**Supplementary Figure 1** | Statistics on the lengths of CDR3β and epitopes in existing databases. **a.** The distribution of sequence lengths of CDR3β shows that the majority of CDR3β sequences consist of 10 to 20 amino acid residues. **b.** The distribution of sequence lengths of epitopes reveals that most antigenic epitopes comprise 8 to 12 amino acid residues. **c.** The distribution of sequence length combinations of CDR3β-epitope pairs in public databases is depicted in the graph. Each dot on the graph represents a specific sequence length combination, where the x-axis corresponds to the sequence length of CDR3β in a CDR3β-epitope pair, and the y-axis corresponds to the sequence length of the epitope in the same pair. Notably, the combination of 15 amino acid residues for CDR3β and 9 amino acid residues for the epitope exhibited the highest number of samples.

#### Supplementary Figures

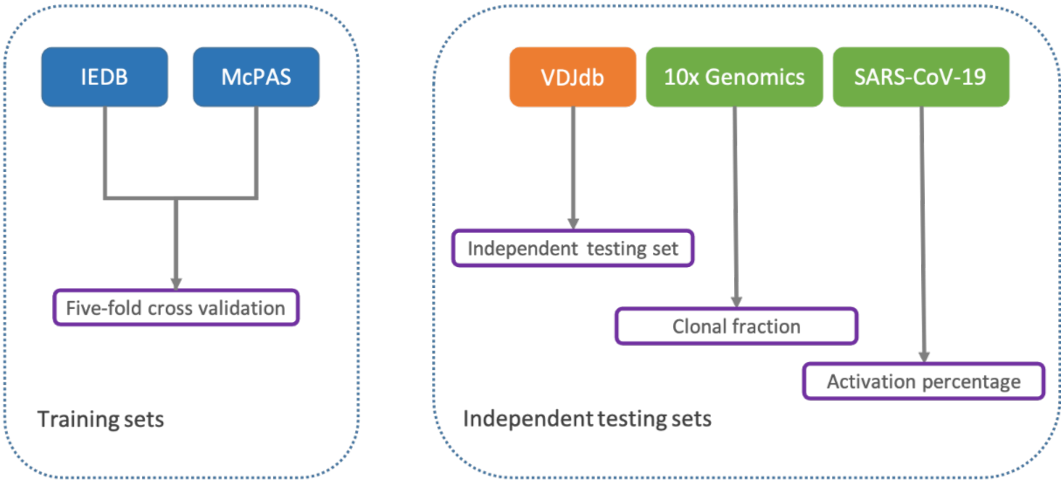


**Supplementary Figure 2** | The schematic diagram illustrates the datasets utilized for model training. Samples from IEDB and McPAS were utilized as the training set, while samples from VDJdb served as an independent test set. The model was further validated based on clonal fraction using samples from 10x Genomics and activation percentage using samples from SARS-CoV-2.

#### Supplementary Figures

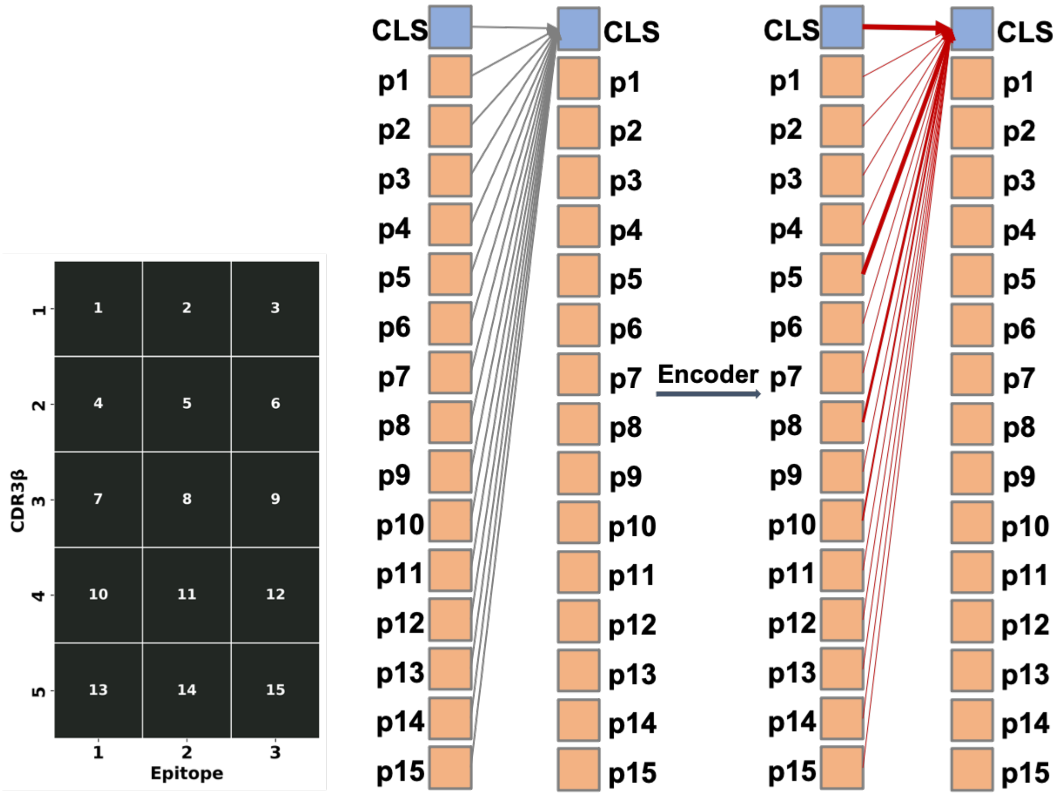


**Supplementary Figure 3 | Illustration of how the self-attention mechanism influences prediction**. The patches numbered in the left panel correspond to the tokens numbered in the right panel. In the right panel, the lines connecting any two tokens represent correlation. Before the Encoder module, the scoring between CLS tokens and other tokens exhibits randomness (indicated by grey lines), while after the Encoder module, the scoring between CLS tokens and other tokens becomes more varied, with some scores being high (represented by thick red lines) and others being low (represented by thin red lines). The thickness of the lines reflects the strength of the score between the two tokens, where thicker lines signify higher scores and thinner lines represent lower scores.

#### Supplementary Figures

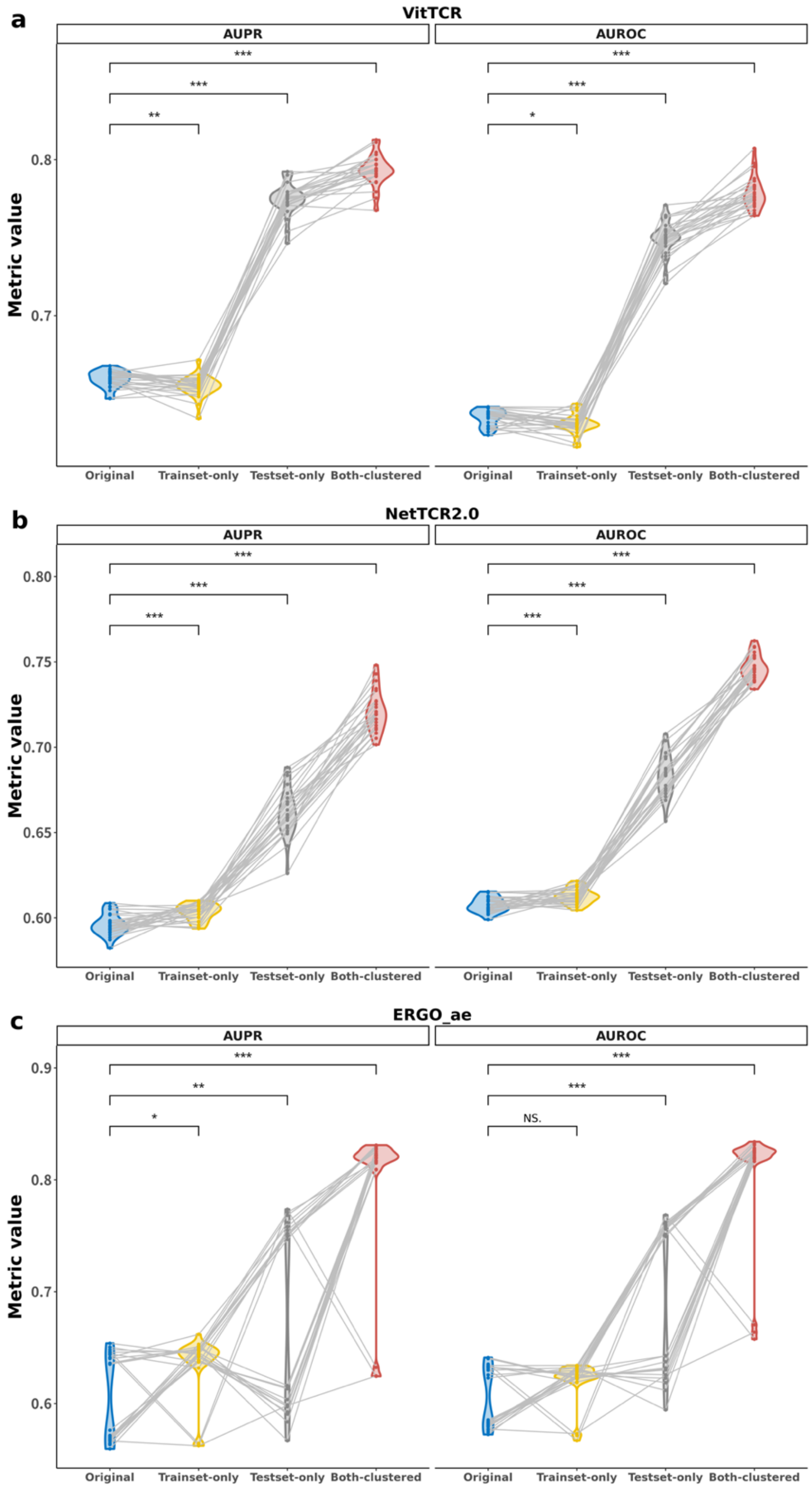


**Supplementary Figure 4 | The influence of cluster-based filtering on model performance in terms of AUROC and AUPR**. The AUROC and AUPR of the three models were compared across the four different dataset settings. Five repeated fivefold cross validations were conducted under four settings, and a dot in this figure represents a fold replicate. **a**. Performance of VitTCR. **b**. Performance of NetTCR-2.0. **c**. Performance of ERGO_AE.

#### Supplementary Figures

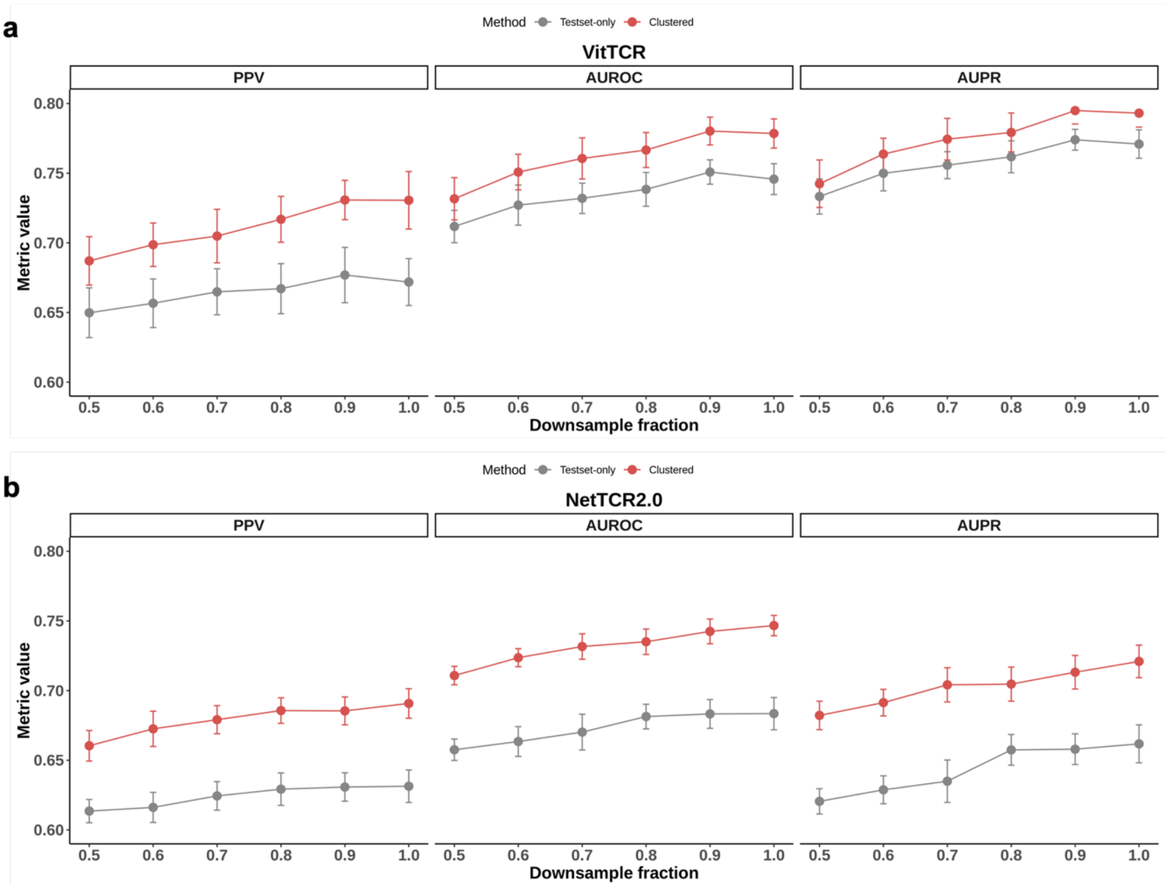


**Supplementary Figure 5 | The impact of cluster-based filtering on model performance in relation to its dependence on the data size**. We performed downsampling on the training set, varying the downsampling ratios from 0.5 to 0.9 in increments of 0.1. For clarity, we focused on two specific settings: "Testset-only" and "Clustered". The two settings utilized the same independent test set while employing different training sets, one with clustering-based filtering and the other without. The downsampling analysis indicates that cluster-based filtering has the potential to decrease the reliance of the model on the size of the dataset.

#### Supplementary Figures

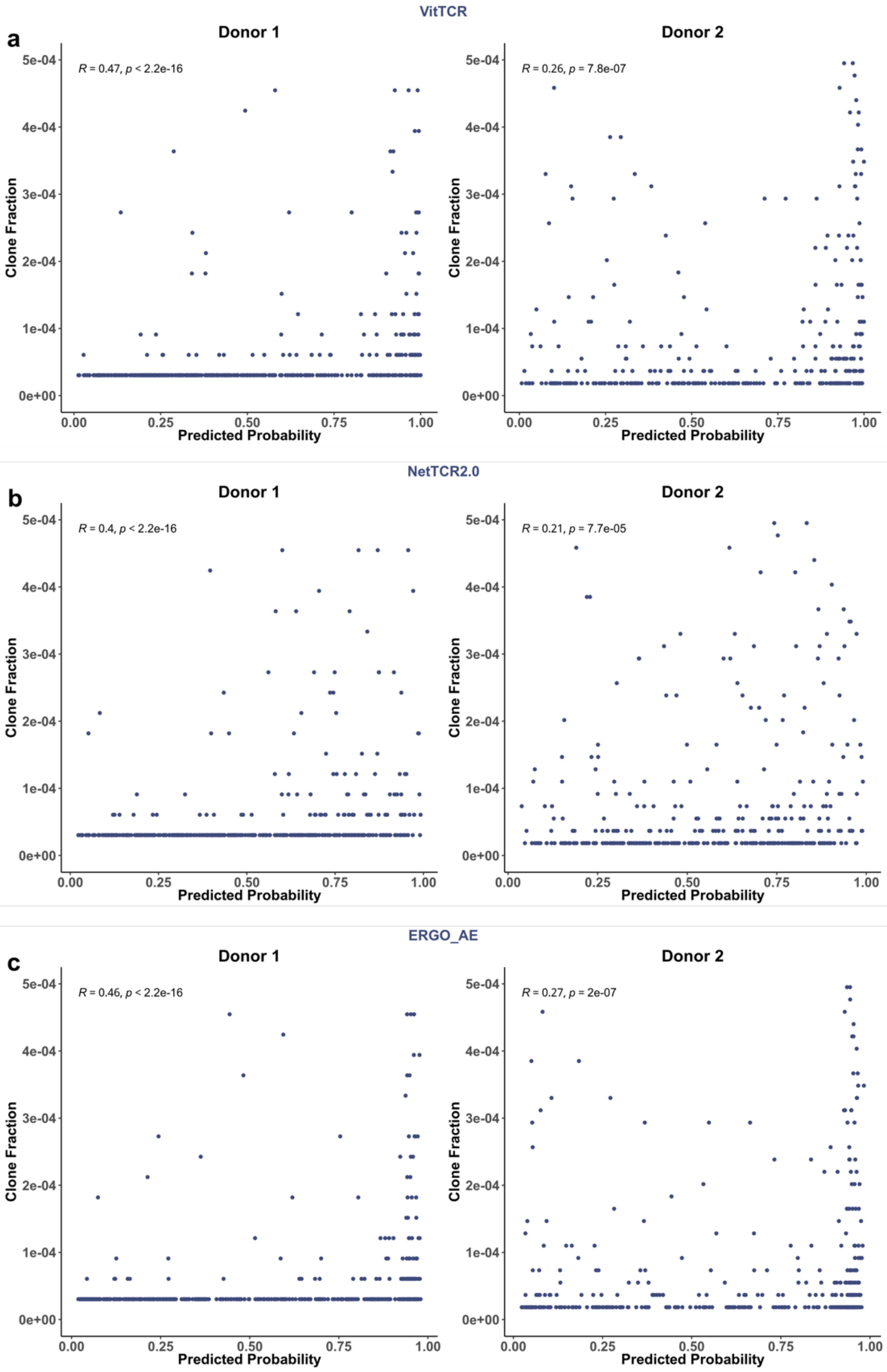


**Supplementary Figure 6 | Influences of cluster-based filtering on model performances**. A dot in the scatter plot represents a TCR clonotype. The x-axis representing the mean of predicted probabilities and the y-axis representing the clone fraction. The Spearman correlation coefficient (R) and significance (p) are also indicated in figure. **a**. Performance of VitTCR. **b**. Performance of NetTCR-2.0. **c**. Performance of ERGO_AE.

### Tables

**Table 1.** The Spearman correlation coefficients and p values between the mean of predicted probabilities and clone fractions for different methods.

| Model | Donor 1 | | Donor 2 | |
|---|---|---|---|---|
| | correlation coefficient | p value | correlation coefficient | p value |
| VitTCR | 0.47 | 2.2e-16 | 0.26 | 7.8e-07 |
| NetTCR-2.0 | 0.40 | 2.2e-16 | 0.21 | 7.e7-05 |
| ERGO_AE | 0.46 | 2.2e-16 | 0.27 | 2e-07 |

#### Supplementary Tables

**Supplementary table 1.** Comparison of clustering before and after across databases (HLA-A*02:01-related data).

| Database | Pair | | CDR3β | | Epitope | |
|---|---|---|---|---|---|---|
| | Before | After | Before | After | Before | After |
| IEDB | 18408 | 9681 | 17760 | 9408 | 321 | 189 |
| McPAS | 1514 | 534 | 1399 | 519 | 23 | 18 |
| VDJdb | 6986 | 2316 | 6791 | 2245 | 552 | 251 |

#### Supplementary Tables

**Supplementary table 2.** Summary of the interactions of 83 TCR-pMHC complexes in the PDB database.

| ID | CDR3β | Epitope | pos_ CDR3β | pos_Epitope |
|---|---|---|---|---|
| 6R2L | CASSPLDVSISSYNEQFF | SLSKILDTV | 10 | 8 |
| 6R2L | CASSPLDVSISSYNEQFF | SLSKILDTV | 12 | 7 |
| 3O4L | CSARDGTGNGYTF | GLCTLVAML | 7 | 8 |
| 2NX5 | CATGTGDSNQPQHF | EPLPQGQLTAY | 5 | 7 |
| 2UWE | CASSDWVSYEQYF | ALWGFFPVL | 0 | 0 |
| 5NME | CASSDTVSYEQYF | SLYNTVATL | 9 | 6 |
| 5NME | CASSDTVSYEQYF | SLYNTVATL | 6 | 8 |
| 3UTT | CASSLWEKLAKNIQYF | ALWGPDPAAA | 6 | 6 |
| 5TEZ | CASSLLGGWSEAFF | GILGFVFTL | 9 | 6 |
| 2VLK | CASSSRSSYEQYF | GILGFVFTL | 7 | 6 |
| 2VLJ | CASSSRSSYEQYF | GILGFVFTL | 7 | 6 |
| 2VLR | CASSSRASYEQYF | GILGFVFTL | 0 | 0 |
| 5JHD | CASSIGVYGYTF | GILGFVFTL | 0 | 0 |
| 5E6I | CASSLIYPGELFF | GILGFVFTL | 0 | 0 |
| 5EUO | CASSIRSSYEQYF | GILGFVFTL | 7 | 6 |
| 5ISZ | CASSIFGQREQYF | GILGFVFTL | 0 | 0 |
| 1OGA | CASSSRSSYEQYF | GILGFVFTL | 7 | 6 |
| 5HHM | CASSSRSSYEQYF | GILGLVFTL | 7 | 6 |
| 5HHO | CASSIRSSYEQYF | GILEFVFTL | 7 | 6 |
| 3HG1 | CAWSETGLGTGELFF | ELAGIGILTV | 8 | 7 |
| 3QDG | CASSLSFGTEAFF | ELAGIGILTV | 6 | 7 |
| 4JFF | CAWSETGLGMGGWQF | ELAGIGILTV | 8 | 7 |
| 4JFF | CAWSETGLGMGGWQF | ELAGIGILTV | 6 | 9 |
| 4L3E | CASSWSFGTEAFF | ELAGIGILTV | 0 | 0 |
| 5E9D | CASRPGWMAGGVELYF | ELAGIGILTV | 8 | 9 |
| 5NHT | CASSQGLAGAGELFF | ELAGIGILTV | 0 | 0 |
| 5NQK | CASSQGLAGAGELFF | ELAGIGILTV | 0 | 0 |
| 6DKP | CASSWSFGTEAFF | ELAGIGILTV | 6 | 7 |
| 4QOK | CAWSETGLGTGELFF | EAAGIGILTV | 8 | 7 |
| 6EQA | CAWSETGLGTGELFF | EAAGIGILTV | 8 | 6 |
| 6EQA | CAWSETGLGTGELFF | EAAGIGILTV | 8 | 7 |
| 6EQB | CAWSETGLGMGGWQF | AAGIGILTV | 8 | 6 |
| 3QEQ | CAISEVGVGQPQHF | AAGIGILTV | 8 | 6 |
| 3QEQ | CAISEVGVGQPQHF | AAGIGILTV | 6 | 8 |

| | | | | |
|---|---|---|---|---|
| 3QEQ | CAISEVGVGQPQHF | AAGIGILTV | 10 | 6 |
| 3QDJ | CASSLSFGTEAFF | AAGIGILTV | 0 | 0 |
| 6D78 | CASSWSFGTEAFF | AAGIGILTV | 0 | 0 |
| 5JZI | CASRRGPYEQYF | KLVALGINAV | 0 | 0 |
| 5YXN | CASRRGPYEQYF | KLVALGINAV | 0 | 0 |
| 2BNQ | CASSYVGNTGELFF | SLLMWITQV | 8 | 7 |
| 2BNQ | CASSYVGNTGELFF | SLLMWITQV | 6 | 6 |
| 6RP9 | CASSSPGGVSTEAFF | SLLMWITQV | 0 | 0 |
| 6RPB | CASSYLNRDSALDF | SLLMWITQV | 7 | 7 |
| 6RPB | CASSYLNRDSALDF | SLLMWITQV | 6 | 6 |
| 6Q3S | CASSYVGNTGELFF | SLLMWITQV | 8 | 7 |
| 6Q3S | CASSYVGNTGELFF | SLLMWITQV | 6 | 6 |
| 2F53 | CASSYVGNTGELFF | SLLMWITQC | 6 | 6 |
| 2F53 | CASSYVGNTGELFF | SLLMWITQC | 8 | 7 |
| 2F54 | CASSYVGNTGELFF | SLLMWITQC | 6 | 6 |
| 2F54 | CASSYVGNTGELFF | SLLMWITQC | 8 | 7 |
| 2PYE | CASSYLGNTGELFF | SLLMWITQC | 6 | 6 |
| 2PYE | CASSYLGNTGELFF | SLLMWITQC | 8 | 7 |
| 6VRM | CASSEGLWQVGDEQYF | HMTEVVRHC | 10 | 7 |
| 6VRM | CASSEGLWQVGDEQYF | HMTEVVRHC | 11 | 7 |
| 6VRM | CASSEGLWQVGDEQYF | HMTEVVRHC | 13 | 7 |
| 6VRM | CASSEGLWQVGDEQYF | HMTEVVRHC | 14 | 8 |
| 6VRM | CASSEGLWQVGDEQYF | HMTEVVRHC | 10 | 6 |
| 6VRN | CAISELVTGDSPLHF | HMTEVVRHC | 6 | 7 |
| 6VRN | CAISELVTGDSPLHF | HMTEVVRHC | 7 | 7 |
| 6VQO | CASSIQQGADTQYF | HMTEVVRHC | 6 | 7 |
| 6VQO | CASSIQQGADTQYF | HMTEVVRHC | 10 | 7 |
| 6VQO | CASSIQQGADTQYF | HMTEVVRHC | 6 | 8 |
| 6VQO | CASSIQQGADTQYF | HMTEVVRHC | 7 | 6 |
| 7RM4 | CASSLDPGDTGELFF | HMTEVVRHC | 0 | 0 |
| 6VMA | CASSMGGTYEQYF | ITDQVPFSV | 0 | 0 |
| 6VM7 | CASSITLSSYNEQFF | ITDQVPFSV | 0 | 0 |
| 3GSN | CASSPVTGGIYGYTF | NLVPMVATV | 7 | 8 |
| 5D2L | CASSQTQLWETQYF | NLVPMVATV | 7 | 8 |
| 5D2N | CASSLAPGTTNEKLFF | NLVPMVATV | 9 | 5 |
| 5D2N | CASSLAPGTTNEKLFF | NLVPMVATV | 9 | 6 |
| 7N1F | CASSPDIEQYF | YLQPRTFLL | 6 | 5 |

| | | | | |
|---|---|---|---|---|
| 7RTR | CASSPDIEQYF | YLQPRTFLL | 6 | 5 |
| 7PBE | CASSSANSGELFF | YLQPRTFLL | 6 | 5 |
| 7PBE | CASSSANSGELFF | YLQPRTFLL | 7 | 5 |
| 7PBE | CASSSANSGELFF | YLQPRTFLL | 8 | 5 |
| 7PBE | CASSSANSGELFF | YLQPRTFLL | 7 | 8 |
| 5C07 | CASSLWEKLAKNIQYF | YQFGPDFPIA | 6 | 6 |
| 5C0A | CASSLWEKLAKNIQYF | MVWGPDPLYV | 6 | 6 |
| 1BD2 | CASSYPGGGFYEQYF | LLFGYPVYV | 7 | 8 |
| 1AO7 | CASRPGLAGGRPEQYF | LLFGYPVYV | 7 | 8 |
| 1AO7 | CASRPGLAGGRPEQYF | LLFGYPVYV | 4 | 5 |
| 4FTV | CASRPGLMSAQPEQYF | LLFGYPVYV | 4 | 5 |
| 4FTV | CASRPGLMSAQPEQYF | LLFGYPVYV | 7 | 8 |
| 4MNQ | CASSYQGTEAFF | ILAKFLHWL | 6 | 8 |
| 5W1V | CASSANPGDSSNEKLFF | VMAPRTLIL | 0 | 0 |
| 6AMU | CASSLSFGTEAFF | MMWDRGLGMM | 6 | 6 |
| 6AMU | CASSLSFGTEAFF | MMWDRGLGMM | 7 | 6 |
| 3H9S | CASRPGLAGGRPEQYF | MLWGYLQYV | 10 | 5 |
| 5C0B | CASSLWEKLAKNIQYF | RQFGPDFPTI | 6 | 6 |
| 5NMG | CASSDTVSYEQYF | SLFNTIAVL | 9 | 6 |
| 5NMG | CASSDTVSYEQYF | SLFNTIAVL | 6 | 8 |
| 3PWP | CASRPGLAGGRPEQYF | LGYGFVNYI | 0 | 0 |
| 1QRN | CASRPGLAGGRPEQYF | LLFGYAVYV | 7 | 8 |
| 6TRO | CASSFLMTSGDPYEQYF | GVYDGREHTV | 13 | 4 |
| 6TRO | CASSFLMTSGDPYEQYF | GVYDGREHTV | 7 | 7 |
| 6TRO | CASSFLMTSGDPYEQYF | GVYDGREHTV | 9 | 6 |
| 5C0C | CASSLWEKLAKNIQYF | RQFGPDWIVA | 6 | 6 |
| 7NDT | CASSQDRDTQYF | VMAPRTLI | 8 | 5 |
| 7NDT | CASSQDRDTQYF | VMAPRTLI | 6 | 5 |
| 7NDT | CASSQDRDTQYF | VMAPRTLI | 7 | 6 |
| 2ESV | CASSQDRDTQYF | VMAPRTLIL | 7 | 6 |
| 2ESV | CASSQDRDTQYF | VMAPRTLIL | 8 | 5 |
| 2ESV | CASSQDRDTQYF | VMAPRTLIL | 6 | 5 |
| 5EU6 | CASSFIGGTDTQYF | YLEPGPVTV | 6 | 7 |
| 5HYJ | CASSLWEKLAKNIQYF | AQWGPDPAAA | 6 | 6 |
| 5NMF | CASSDTVSYEQYF | SLYNTIATL | 9 | 6 |
| 5NMF | CASSDTVSYEQYF | SLYNTIATL | 6 | 8 |
| 7N1E | CASSLGGAGGADTQYF | RLQSLQTYV | 7 | 8 |

| | | | | |
|---|---|---|---|---|
| 2GJ6 | CASRPGLAGGRPEQYF | LLFGKPVYV | 6 | 5 |
| 2GJ6 | CASRPGLAGGRPEQYF | LLFGKPVYV | 7 | 8 |
| 3QFJ | CASRPGLAGGRPEQYF | LLFGFPVYV | 7 | 8 |
| 5C08 | CASSLWEKLAKNIQYF | RQWGPDPAAV | 6 | 6 |
| 5C09 | CASSLWEKLAKNIQYF | YLGGPDFPTI | 6 | 6 |
| 1QSE | CASRPGLAGGRPEQYF | LLFGYPRYV | 6 | 7 |
| 1QSE | CASRPGLAGGRPEQYF | LLFGYPRYV | 10 | 7 |
| 1QSE | CASRPGLAGGRPEQYF | LLFGYPRYV | 7 | 8 |
| 4EUP | CASSFLGTGVEQYF | ALGIGILTV | 6 | 8 |
| 4EUP | CASSFLGTGVEQYF | ALGIGILTV | 8 | 3 |
| 4EUP | CASSFLGTGVEQYF | ALGIGILTV | 8 | 6 |

The table includes the “ID” column, which corresponds to the identification number of the TCR-pMHC complex in the PDB (Protein Data Bank) database. Each row in the table provides the amino acid sequences of CDR3β (found in the “CDR3β” column) and the epitope (listed in the “Epitope” column) within a TCR-pMHC complex. Furthermore, the positions of amino acids involved in interactions are recorded in the “pos_CDR3β” and “pos_Epitope” columns. Certain TCR-pMHC complexes, such as 6R2L, are represented by two rows in the table, indicating the presence of two pairs of interacting amino acids within the complex. Conversely, some TCR-pMHC complexes, such as 2UWE, have “zero” values in both the “pos_CDR3β” and “pos_Epitope” columns. Importantly, this does not imply that the CDR3β of the complex does not recognize its cognate epitope. Rather, it suggests that the interactions of the complex might not have been captured during the structural analysis using PyMOL. To ensure the accuracy of our analysis regarding the positional bias weight matrix, we excluded samples in which no interactions were observed in PyMOL. We accomplished this by removing samples with “ zero” values in both the “pos_CDR3β” and “pos_Epitope” columns. As a result, we retained 101 pairs of amino acids derived from 61 TCR-pMHC complexes that exhibited interactions. Each amino acid pair comprised one amino acid from the CDR3β region and another from the epitope.
